# Cell atlases and the Developmental Foundations of the Phenotype

**DOI:** 10.1101/2025.01.20.633901

**Authors:** Alicia Lou, Mónica Chagoyen, Juan F Poyatos

## Abstract

It is widely acknowledged that development shapes phenotypes, yet the extent to which genes with similar expression patterns during development lead to equivalent organismal phenotypes when mutated remains unclear. Here, we propose addressing this issue, which we term the 𝒟evelopment–to– 𝒫henotype, or 𝒟–𝒫, rule, by leveraging single-cell gene expression atlases and phenotypic ontologies, using *Caenorhabditis elegans* as a model system. This framework quantifies the proportionality between developmental expression and phenotypic similarities, demonstrating that the relationship holds on average. Genes that strongly fulfill the rule exhibit broad “housekeeping” expression and are associated with systemic phenotypes, whereas weak similarities correspond to specific expression patterns and specialized phenotypes. Deviations from the 𝒟–𝒫 rule provide insights into developmental divergence and phenotypic degeneracy, highlighting genes with narrow functional roles but systemic phenotypic impact. Furthermore, genes that closely adhere to the rule exhibit the highest pleiotropic impact on organismal traits. Our analysis also identifies cell types, such as ASK neurons, as key mediators of phenotype-specific gene contributions, exemplified by their association with chemosensory behavior and chemotaxis. These findings validate the 𝒟–𝒫 rule and underscore the role of cells as critical mediators of the genotypephenotype map, offering a unified framework to understand the developmental origins of phenotypic complexity.

## Introduction

Cutting-edge sequencing technologies now enable precise tracking of gene activity at single-cell resolution during embryonic development, surpassing earlier methods that could only assess embryos in bulk, e.g., (Hashimshony et al. [2015]). This advancement is driving the creation of cell atlases –comprehensive maps of cell states and types– providing new insights into developmental dynamics across a wide range of organisms, including zebrafish (Wagner et al. [2018]), western claw-toed frog (Briggs et al. [2018]), nematode (Packer et al. [2019]), fruit fly (Calderon et al. [2022]), or mouse (Qiu et al. [2024]).

With the availability of these atlases, numerous issues can now be addressed. For example, they can be applied to better understand the relationship between gene expression, cellular functions, and cell differentiation (Green et al. [2024], Liu et al. [2024]), or to provide a comprehensive gene expression map within organoids, facilitating validation against *in vivo* counterparts (Watanabe et al. [2017]). At the same time, these data enable a reexamination of fundamental biological questions, such as uncovering the enhancer logic underlying cell identity (Bravo González-Blas et al. [2023]), determining the proper approaches to defining and organizing cell states and types (Domcke and Shendure [2023]), and exploring the role of cellular plasticity in developmental robustness (Xiao et al. [2022]), among others.

This study falls into the last type of questions. Our goal is to leverage data from embryonic cell atlases to deepen our understanding of how development shapes phenotypes. Although this question is necessarily broad –development both constrains (Alberch [1980]) and diversifies (West-Eberhard [2003]) phenotypes– we simplify it to a *quantitative* evaluation of a fundamental “rule” that addresses how the sequence of developmental events influence phenotypic outcomes. Specifically, we ask: do genes with similar developmental expression patterns consistently lead to equivalent phenotypic outcomes at the organismal level? While this is a simplification of development’s full complexity within the genotype-to-phenotype (GP) map, it offers an approach we can rigorously evaluate. We refer to the validation of this hypothesis as the 𝒟evelopment–to–𝒫henotype, or 𝒟– 𝒫, *rule*.

However, evaluating this rule presents several challenges. First, accurately computing the developmental details within cellular atlases requires integrating how genes are activated or silenced at specific stages and in particular cell types. Second, while *cellular* phenotypes can be obtained on a relatively large scale, e.g., (Green et al. [2024]), capturing *organismal* phenotypes poses a greater difficulty. These two obstacles are addressed using *Caenorhabditis elegans* as a model system.

The first goal is to formalize the developmental information encoded in cell atlases that map the organism’s development. To achieve this, a unified developmental signature is proposed, integrating both embryonic time and cell type, thereby enabling the characterization of each gene’s activity within this context. This effort utilizes a recent dataset where most cells have been annotated with a cell type or lineage across stages of embryogenesis, primarily from mid-gastrulation to terminal differentiation (Packer et al. [2019]). Our approach will focus exclusively on annotated cell types. For the second challenge, the Worm Phenotype Ontology (WPO) –a hierarchically structured, controlled vocabulary for standardizing phenotypes (Schindelman et al. [2011])– is employed to represent a phenotypic space where each gene is associated with a defined subset of phenotypes. Intrinsic redundancies in the ontology are addressed using non-negative matrix factorization (NMF) (Lee and Seung [1999]).

Armed with these datasets, we aim to address three primary questions. First, we introduce an *average* developmental and phenotypic similarity score for each gene to quantify how closely its developmental or phenotypic patterns resemble those of others. Genes with higher developmental similarity also tend to have higher phenotypic similarity, consistent with the 𝒟–𝒫 rule. However, some genes deviate from this pattern, prompting us to investigate the mechanisms underlying both phenotypic degeneracy (Edelman and Gally [2001]) and developmental divergence (Carroll et al. [2005]). Next, we examine how varying levels of developmental specificity correlate with a gene’s impact on phenotypes, i.e., pleiotropy (Zhang [2023]). This we do by distinguishing a set of pleiotropic genes. We also identify specific developmental “coordinates” where pleiotropic genes are dominantly expressed. These genes show enrichment particularly at early stages. Finally, we explore whether certain cell types act as key mediators of phenotypic outcomes, referring to this as a *coarse* interpretation of the rule. This work demonstrates how integrating cell atlases and phenotypic ontologies reveals the extent to which developmental processes shape phenotypic outcomes, while also providing a complementary perspective on pleiotropy and the role of cells as mediators in the GP map.

## Results

### A simple rule within the genotype-phenotype map

To evaluate how similar developmental gene expression trajectories lead to comparable organismal phenotypes, the 𝒟–𝒫 rule, we establish two quantitative spaces: a developmental space and a phenotypic space. The developmental space reflects the distribution of gene expression across cell types (Table S1) and embryonic stages, while the phenotypic space captures the impact of gene deactivation on organismal phenotypes (Figure 1A). Each gene is assigned a pair of vectors, one from each space, enabling direct comparisons between developmental patterns and phenotypic outcomes (Methods, extended analysis in Supplement, Figs. S1-S6); 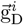 represents the fraction of cells sampled at a given developmental coordinate in which the corresponding gene *i* is activated, while 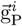 quantifies the gene *i*’s contribution to an NMF-derived phenotype space.

**Figure 1.**
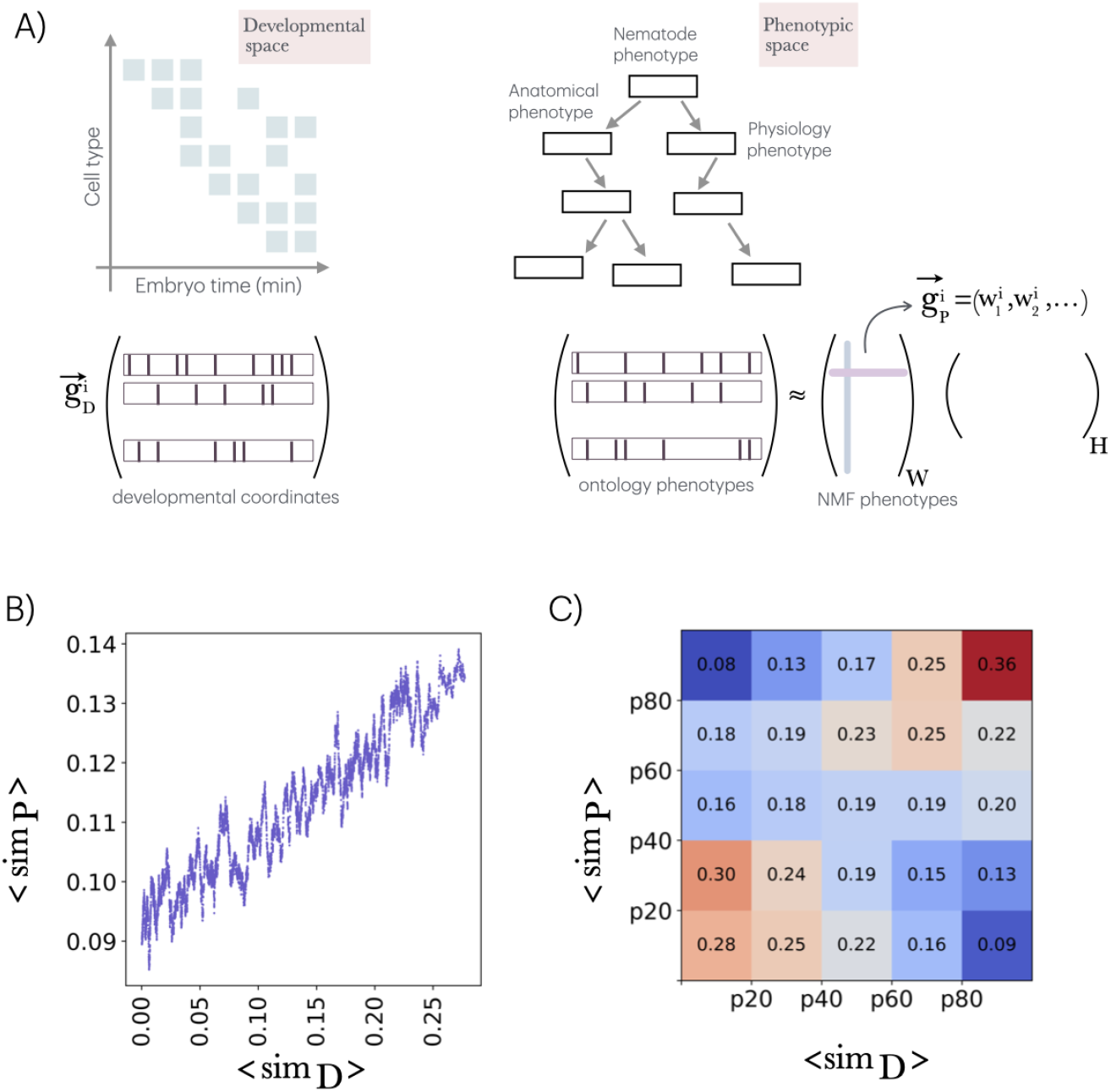
Developmental and phenotypic spaces and the 𝒟–𝒫 rule. A) Developmental and phenotypic spaces are characterized using data from the single-cell atlas and the worm ontology, respectively. For each gene *i*, the relative number of cells in which it is active within the sampled set at each specific developmental coordinate (cell type and embryonic time) constitutes the elements of its associated developmental vector, 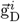. Collectively, these vectors define the developmental matrix. Similarly, a phenotypic (binary) matrix is constructed by associating genes with their linked phenotypes in the ontology. This matrix is decomposed using nonnegative matrix factorization, NMF (into a *features* matrix W and a *coefficients* matrix H). In this decomposition, the gene phenotypic vector 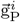 is represented by the rows of W (highlighted in pink), while each column (in blue) indicates the contribution of each gene to a specific NMF-derived phenotype. B) By defining and average (median) similarity score for each gene with respect to the rest, we observed a proportionality between ⟨*sim*_D_⟩ and ⟨*sim*_P_⟩ what highlights the 𝒟–𝒫 rule: “similar developmental trajectories lead to similar phenotypes”. Plot is a slidding window of the full data (windows size = 100). C) For each set of genes with an average similarity ⟨*sim*_D⟩_ within specific percentiles (e.g., p20 indicating less or equal to the 20th percentile, and so on), the percentage of genes belonging to different phenotypic percentile categories is presented (each column sums to 1). Deviations from the expected percentage in development and phenotype suggest a departure from the rule. See main text for further details.

Using this information, we compute two similarity matrices –one for development and another for phenotype– where each entry represents the pairwise similarity between any two genes in the respective space [Methods, Tables S2-S3 include similarity values and gene ontology (GO) analysis as functions of these similarities]. We then simplified these matrices by computing an average developmental similarity ⟨*sim*_D_⟩ and phenotypic similarity ⟨*sim*_P_⟩ for each gene, which represent the median of all its associated pairwise values. If the 𝒟–𝒫rule holds, one should observe a proportional relationship between these two measures. We confirm this relationship (Figure 1B; Supplement, Figs. S7-S8). To further examine the strength of this rule, genes were categorized into distinct ranges of the corresponding ⟨*sim*_D_⟩ and ⟨*sim*_P_⟩ distributions (below the 20th percentile, between the 20th and 40th percentiles, and so on). The frequency of the relevant phenotypic categories for each developmental class was computed. According to a *strict* 𝒟–𝒫 rule, phenotypic categories should correspond one-to-one with developmental ones. We do observe a tendency for genes to be part of equivalent classes; however, we also noted deviations, with some genes showing equivalent developmental patterns but differing phenotypic ones, and vice versa (Figure 1C). These deviations will be studied next.

### Exceptions to the 𝒟–𝒫 rule

To analyze discrepancies in the rule, we first fit ⟨*sim*_P_⟩ against ⟨*sim*_D_⟩ using Loess regression and calculated the residuals for each data point, i.e., deviations from the fit (Supplement, Figs. S9-S10).

This allowed us to define a group that follows the rule with high ⟨*sim*_D_⟩ and ⟨*sim*_P_⟩ (that we termed `D-P` genes) and identify two main deviations (`D-p` and `d-P` genes, Figure S9A).

Figure 2 shows representative gene expression patterns for each class, with 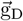 simplified by averaging across cell types at each embryonic time point (Methods). Developmental *divergent* genes (`D-p`, which are developmentally similar but phenotypically distinct) are expressed across most cell types and embryonic stages, similar to `D-P` genes (Figure S9B). However, their phenotypic roles differ: while `D-P` genes are associated with systemic and general phenotypes, `D-p` genes are linked to a specific subset of enriched phenotypes (Table S4). This distinction suggests that `D-P` genes act as housekeeping genes essential for fundamental processes across all cell types, with their disruption leading to systemic organismal failure. In contrast, `D-p` genes, although also housekeeping genes, play critical roles only in specific cellular contexts. Their loss does not cause a complete systemic failure but results in localized dysfunctions tied to the cell types where their function is indispensable.

**Figure 2.**
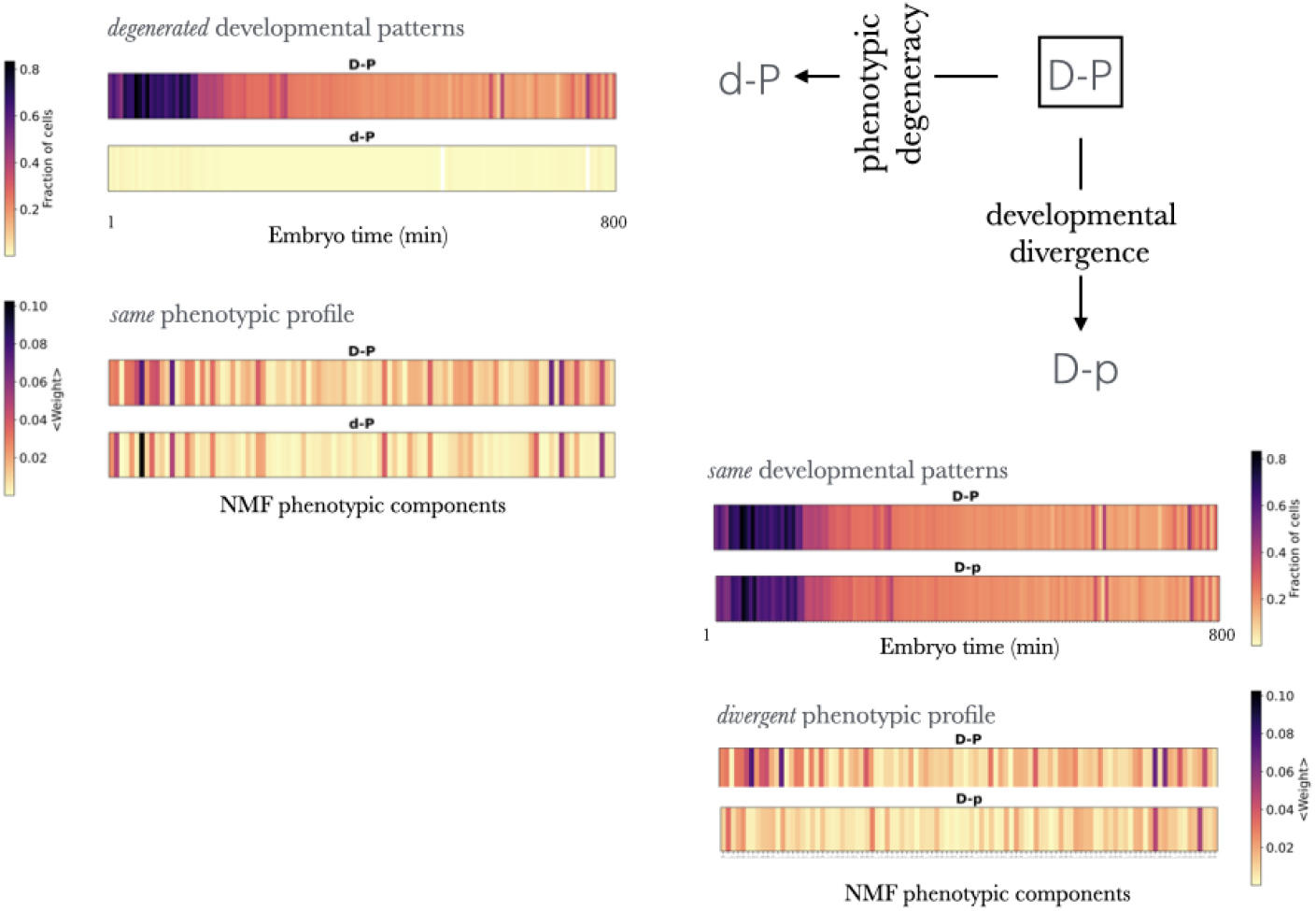
Deviation from the D-P rule. We consider the subclass of genes that fulfill the – rule 𝒟–𝒫 and show strong similarity, denoted `D-P`, to examine two deviations: phenotypic *degeneracy* or developmental *divergence*. In the case of divergence, denoted as `D-p`, deviations occur when genes show similar developmental patterns to the `D-P` class but differ in their phenotypic outcomes (bottom right of the figure). These patterns are represented as barcodes, reflecting the characteristic gene expression of a gene within the corresponding class (averaged across the cell types at each embryonic stage) or its contribution to phenotypic components. In the case of degeneracy, denoted as `d-P`, deviations occur when genes display different developmental patterns from the `D-P` class but result in similar phenotypes (top left of the figure). For details, refer to the main text, Methods, Supplement, and Fig. S10.

Phenotypic *degeneracy* is characterized by genes with similar phenotypic profiles arising from multiple developmental patterns (`d-P`). Unlike `D-P` genes, which are housekeeping genes critical for the organism’s overall viability, `d-P` genes exhibit specific developmental profiles, influencing particular cell types or embryonic stages while their associated phenotypes remain systemic and general. Although `d-P` genes are active in a limited number of cell types, their localized functions are vital for organismal survival, with lethality often being one of their enriched phenotypes. See Supplement and Fig. S10 for additional details and a GO analysis revealing additional differences between the three groups (Table S5).

### The 𝒟–𝒫 rule and the pleiotropy of complex phenotypes

Since genes that exhibit strong similarities and align with the rule’s premise impact many phenotypes, we next aimed to examine this more comprehensively using an NMF-derived pleiotropic score, 𝕡. This score quantifies the number of traits influenced by a single gene, identifying pleiotropic genes as those exceeding the 95th percentile of the distribution (Supplement, Fig. S11). NMF’s ability to isolate fundamental phenotypic elements overcomes limitations of traditional pleiotropy measures, which are often constrained by the complexity and interdependence of phenotypes (Zhang [2023], Zou et al. [2008]) (Fig. S12).

We began by first examining the expression patterns of pleiotropic genes. For each gene, we calculated the ratio of cells (regardless of type or embryonic time) in which it is expressed relative to the total number of cells, then averaged these ratios. The results showed that pleiotropic genes are expressed in significantly more cells than expected. In contrast, non-pleiotropic genes (those below the 5th percentile of the distribution, Figure 3A) are expressed in far fewer cells and show no notable enrichment for phenotypes or GO terms. Pleiotropic genes, however, are strongly associated with systemic phenotypes and functions (Table S6-S7).

**Figure 3.**
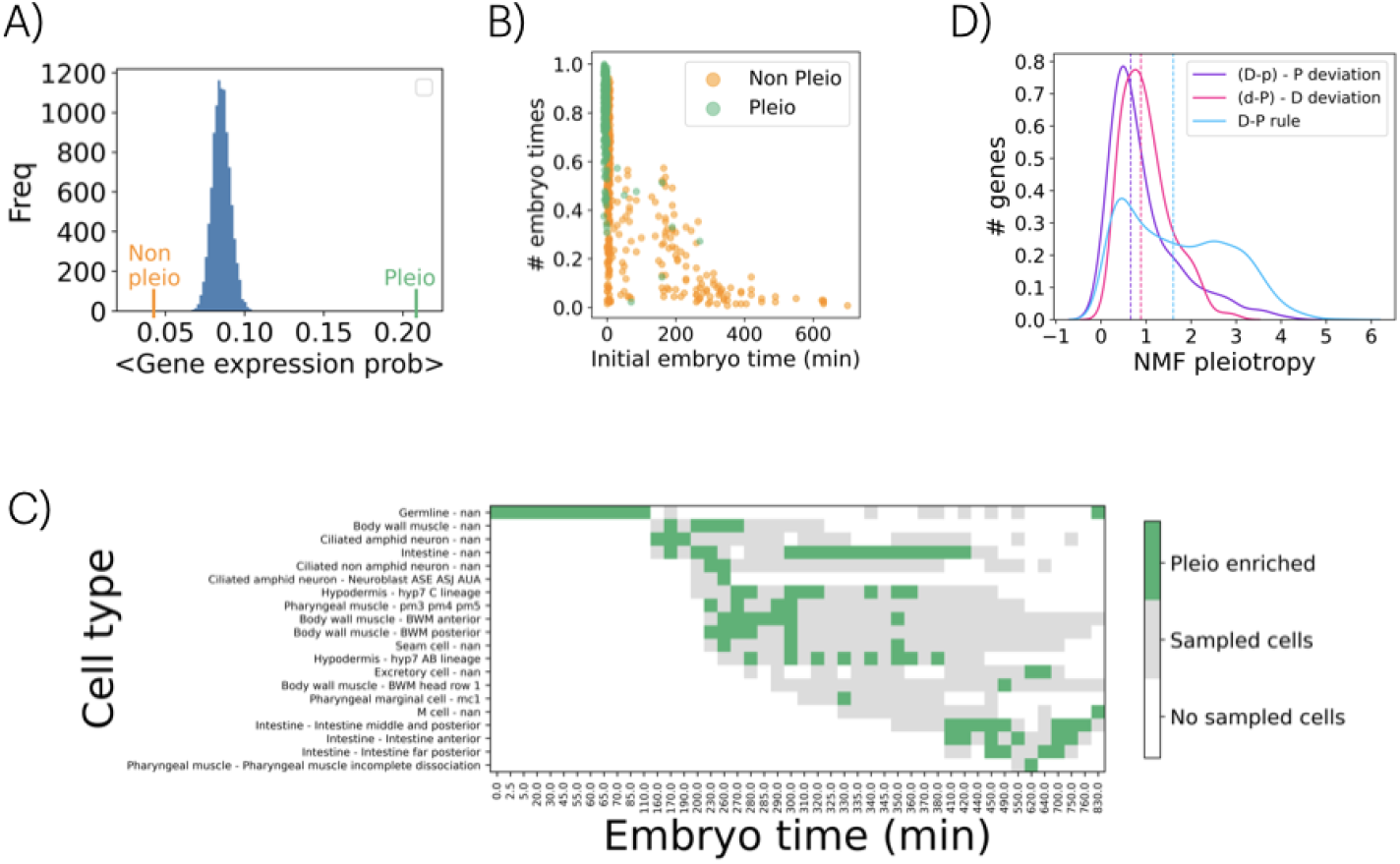
Pleiotropy and the 𝒟–𝒫 rule. A) Average gene expression probability for non-pleiotropic genes (orange, *n* = 418) and pleiotropic genes (green, *n* = 412). Additionally, the distribution of 10,000 averages of gene expression probabilities, calculated by randomly selecting n = 415 genes from the dataset, is shown. B) Each gene of the two sets is shown in an space of initial expression embryo time and fraction of different embryo times in which a genes is expressed. Pleiotropic genes tend to be initially expressed at earlier times in a higher number of stages. C) Distribution of the top 100 pleiotropic genes across developmental coordinates. Enriched regions are highlighted in green, with sampled cell coordinates shown in gray. Note a subgroup associated with specific intestine and pharyngeal cell types, both appearing relatively late in development (lower right). D) Kernel density estimates of pleiotropic score distributions for D-p, d-P, and d-P genes. The *x*-axis represents pleiotropy scores, and the *y*-axis indicates the fraction of genes in each group. Dashed lines show medians, with a mean pleiotropy score of 3.5 for pleiotropic genes.

To incorporate developmental context, we extended the analysis in two ways: by assessing enrichment across embryonic times and by considering the full (cell type, embryonic time) coordinates (Figure 1A). We first computed both the range of embryonic stages each gene is expressed in, as well as the earliest stage of expression. Pleiotropic genes tend to be expressed earlier and across more stages than non-pleiotropic ones (Fig. 3B, Fig. S13). We then analyzed developmental coordinates (cell type, embryo stage) enriched for pleiotropic genes by calculating a *z*-score for expression likelihood at each coordinate (Supplement, Fig. S13D). Figure 3C shows the top 100 coordinates enriched for these genes. There exists again a tendency for pleiotropic genes to be enriched in cell types identified at early developmental stages. Note also a subgroup connected to specific intestine and pharyngeal cell types (lower right in Fig. 3C); both classes appearing relatively late during development.

Moreover, to examine more explicitly the relationship between the 𝒟–𝒫 rule and pleiotropy, we compared the pleiotropy distributions of the three groups of genes (`D-P`, `d-P` and `d-P`) obtained in the previous section depending on their adherence to the rule (Figure 3D). Genes that are strongly similar and fulfill the rule (`D-P` genes) are the ones with the highest pleiotropy values. Moreover, since high ⟨*sim*_P_⟩ values are typically associated with elevated ℙ (Fig. S14), this relationship explains the shift toward higher ℙ observed in `d-P` genes compared to the `D-p` class.

### A coarse interpretation of the 𝒟–𝒫 rule

Finally, we take a coarser yet complementary approach to the 𝒟–𝒫 rule by grouping the developmental space based on cell types. This method allows us to investigate whether genes expressed in specific cell types are strongly linked to particular phenotypes (Figure 4A), offering an alternative perspective for exploring development–phenotype associations.

**Figure 4.**
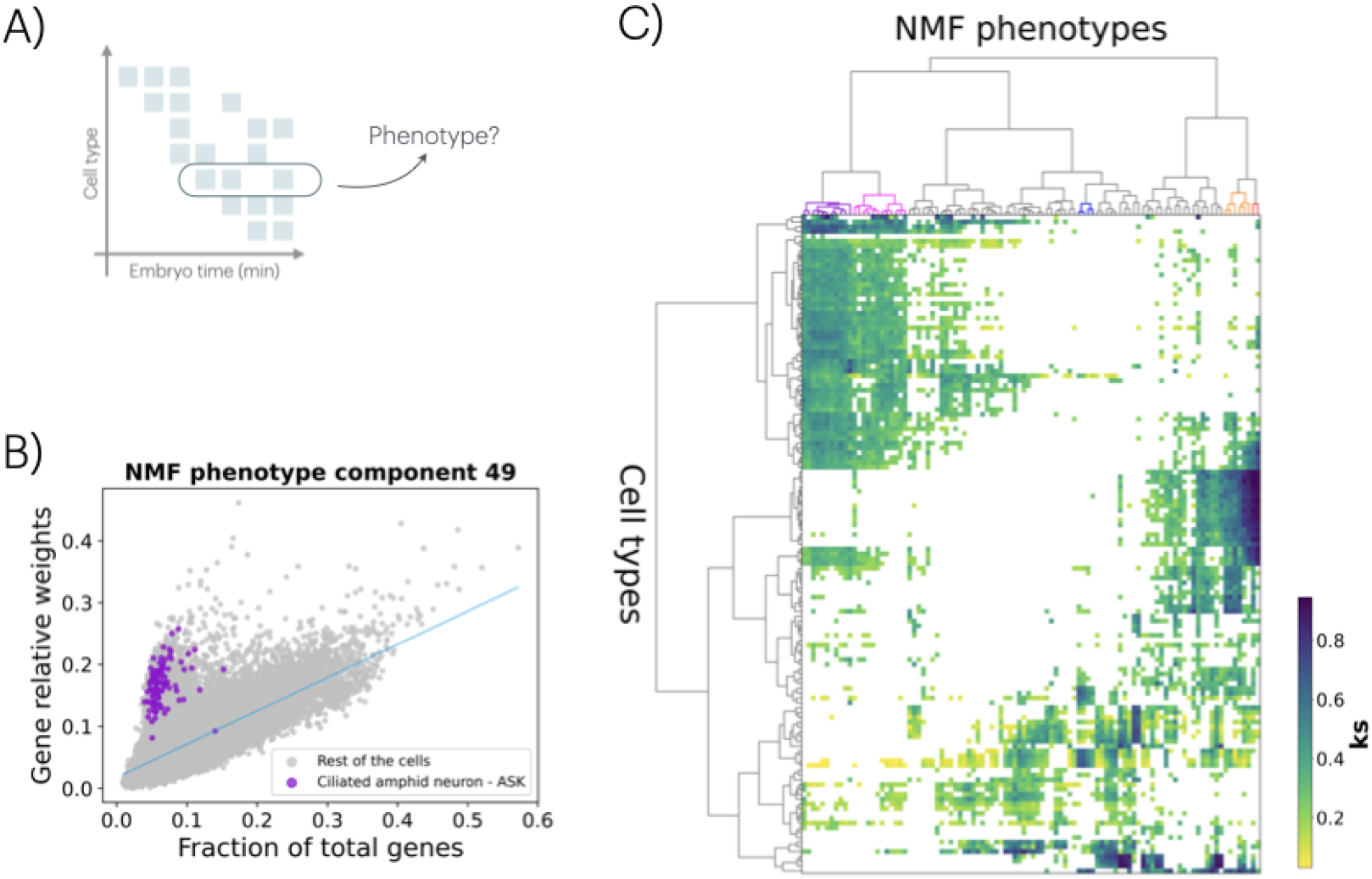
Coarse interpretation of the D-P rule. A) The original developmental space (see Fig. 1A) can be simplified by grouping cells by type to assess how strongly these cell types mediate phenotypes. B) By calculating the fraction of expressed genes in each cell (each cell represented as a dot) and the relative contribution of these genes to a specific NMF-derived phenotype (in this case, component #49), we identify a regression pattern. Cells with larger residuals indicate enrichment for the phenotype, with violet dots corresponding to ASK ciliated amphid neurons. This association is determined by comparing the residual distribution of cells from a specific type to that of all other cells using a Kolmogorov-Smirnov (KS) test. Across all cell types, we find a significant association between ASK neurons and phenotype #49 (KS = 0.94, p-value < 0.0001). C) By analyzing all NMF phenotypes and identifying cell types with significant KS associations, we generate a heatmap linking NMF phenotypes to cell types. The heatmap’s color represents the KS statistic for each association, and all plotted KS values are significant (*p*-value < 0.0001). The *x*-axis lists sorted NMF phenotypes, while the *y*-axis lists cell types, both organized using hierarchical clustering. Dendrograms for each axis are included, grouping related cell types into broader categories, such as precursor and parent cell types, those associated with the nervous system, and others linked to specific anatomical structures (e.g., pharynx, hypodermis, intestine, body wall).

To attain this, we calculate the fraction of genes expressed in a given cell and the relative contribution of these *active* genes to each NMF-derived phenotype (Methods). The results reveal an expected trend in which cells with a greater number of expressed genes typically display higher cumulative weights for the corresponding phenotype (Figure 4B). However, by performing regression on this trend, we identified cells where the expressed genes have a disproportionately strong influence. If these cells belong to a specific cell type, that type can be identified as a “mediator” of the phenotype using a Kolmogorov-Smirnov (KS) statistical test applied to the distribution of the corresponding regression residuals (Methods). For example, Figure 4B highlights the association between the ASK ciliated sensory neuron and NMF phenotype #49 (KS statistic = 0.94; see also Fig. S15 for additional examples). This phenotype is linked to WPO terms like chemosensory behavior, chemotaxis, and odorant response, corroborating the known role of ASK neurons in detecting chemical stimuli and supporting the relevance of the associations and the statistical approach.

Figure 4C extends this analysis, illustrating the connections between all NMF-derived phenotypes and cell types. Each cell type is linked to at least one phenotype and each phenotype is connected to at least one cell type (Fig. S16). Closer examination of the figure highlights that certain cell types have a stronger influence on complex phenotypes, while some phenotypes are less/more closely associated with specific cell types (Supplement, Table S8). Figure 4C also shows distinct cell and phenotype groupings that stand out for having associations with high KS scores: ciliated amphid neurons with chemosensory responses (red in the top dendogram), neurons with acetylcholinesterase inhibitor response (orange), body wall and pharyngeal muscle cells with muscle morphology phenotypes (dark blue), and germline cells with sterile progeny and lethal phenotypes (magenta) or reduced fertility phenotypes (purple). Broader *cell* groupings associated with similar phenotypes are also notable (Fig. S17A), including precursor and parent cells, nervous system-associated cells, and cells linked to specific anatomical structures (e.g., pharynx, hypodermis, intestine, body wall, Table S9), with enrichment of these cell types in the *C. elegans* AB lineage (Fig. S17B, Supplement).

## Discussion

It is commonly proposed that equivalent gene expression during embryogenesis among a set of genes suggests their potential involvement in analogous organismal phenotypes (Carroll et al. [2005]). However, quantitatively testing this hypothesis has been challenging. For instance, we lacked single-cell resolution gene expression measurements during development, and large-scale quantification of organismal phenotypes was similarly constrained. By utilizing cell atlases and phenotypic ontologies, we aim to overcome these limitations.

One approach to leveraging these datasets is by defining a developmental vector and a phenotypic vector for each gene. This quantification allows us to introduce similarity measures –specifically, to assess how similarly any two genes are expressed during development or how similarly they are linked to phenotypes when perturbed. The 𝒟–𝒫 *rule* hypothesizes proportionality between developmental and phenotypic similarity, which our analysis confirms on average (Figure 1). The rule suggests that “house-keeping” activity (i.e, ubiquous expression during development) correlates with broader involvement in common phenotypes, while more specific expression profiles align with specialized phenotypes.

There are also deviations from this rule, two of which we examined (Figure 2). First, genes with high expression similarity across many cell types and developmental stages can exhibit diverse phenotypes (`D-P` *υs*. `D-p` ; recall that capitals denote strong similarity, while lowercase indicates dissimilarity; thus, `D-P` genes exhibit strong developmental and phenotypic similarity, adhering to the rule). Despite their broad expression, `D-p` genes are associated with particular phenotypes, explaining their lower ⟨*sim*_P_⟩ values. A second deviation involves phenotypic degeneracy (`D-P` *υs*. `d-P`). In this case, the expression of certain `d-P` genes at distinct developmental stages can drive broad systemic phenotypic changes. These genes are enriched in only a small number of GO terms, reflecting a general heterogeneity of specific cellular functions crucial for organismal viability, such as nucleosome and chromatin functions (Supplement, Table S5).

That genes that are broadly expressed or serve diverse roles during development might contribute to varied phenotypic effects takes us to the very concept of pleiotropy (Xiao et al. [2022], Zhang [2023], Zou et al. [2008]). To examined this, we introduced a quantitative measure based on adding the scores associated with each gene in the features matrix W obtained by NMF (Lee and Seung [1999]). Pleiotropic genes are expressed in a significantly larger proportion of cells compared to non-pleiotropic genes, as one would anticipate, with broad and early-stage expression being additional features (Figure 3). Notably, this measure correlates with one derived from an experimentally quantified set of cellular phenotypes linked to gene knockouts from *C. elegans* chromosome I expressed during embryogenesis (Xiao et al. [2022]) (Spearman’s ρ = 0.57, p = 3.36×10^−25^, Supplement), reinforcing a view of the cell as a mediator of phenotypes, even when cellular plasticity might partially decouple this association (Xiao et al. [2022]).

These findings corroborate the notion that pleiotropy is linked to genes with foundational roles in early development, which influence diverse downstream processes. Despite their early and broad expression, subsets of pleiotropic genes exhibit late-stage specificity (e.g., intestine and pharyngeal cell types, Figure 3C), indicating that pleiotropy can also manifest in temporally constrained ways. Finally, the highest pleiotropic scores were observed in genes that exhibit strong similarities and fulfill the 𝒟–𝒫 rule (`D-P` class), as high phenotypic similarity correlates with increased pleiotropy (⟨*sim*_P_⟩ scales with ℙ).

In exploring the development-phenotype associations, we grouped cells based on their types and examined their contributions to NMF-derived phenotypes. We identified cell types where those genes expressed during development disproportionately influenced the phenotype. For example, most genes activated in ASK neurons are linked to NMF phenotype #49 (Figure 4), which strongly associates with chemosensory behavior and chemotaxis, aligning with its known role in chemical stimulus detection. This coarse approach enabled us to group phenotypes connected to specific cell types and cell groups associated with similar phenotypes (Figure 4C, Supplement) what validates cells as a “mediator” of the GP map.

Intrinsic features of *C. elegans* development could complicate, but if properly accounted for, potentially broaden the applicability of our approach to the 𝒟–𝒫 rule. These features include the convergence of different lineages to produce the same cell type and the minimal differences in cell expression observed until the final two rounds of cell division (Liu and Murray [2023], Packer et al. [2019]). Nevertheless, this work demonstrates the utility of our simplification in validating the 𝒟–𝒫 rule. Notably, our approach provides a means of assessing the extent to which similarity in phenotypic traits clusters genes whose expression is active in shared developmental processes. This offers a valuable tool for functional genetic profiling, particularly for disease contexts, including rare and poorly understood conditions with unknown genetic bases (Lee et al. [2020]). Collectively, these efforts contribute to unraveling the developmental underpinnings of phenotypes and advancing our understanding of how developmental processes shape organismal traits.

## Materials and methods

### scRNA-seq data

We used the developmental cell atlas of *C. elegans* produced by Packer *et al*. (Packer et al. [2019]). The corresponding data is available at the Gene Expression Omnibus (GEO) repository (www.ncbi.nlm.nih.gov/geo) under accession code GSE126954. This dataset includes transcriptome information for 20,222 genes across 86,024 single cells, represented as unique molecular identifier (UMI) counts. For each cell, developmental timing was estimated by correlating its transcriptome with a bulk RNA-seq time series (Hashimshony et al. [2015]). Additionally, the combination of marker genes, lineage assignments, and developmental timing enabled annotation of the majority of cells by cell type or lineage. (Supplement, Fig. S5)

### Worm Phenotype Ontology data

We downloaded *C. elegans* phenotypes and their gene associations (version WS290) from the Worm Phenotype Ontology (Schindelman et al. [2011]) found in the WormBase (www.wormbase.org/). Gene-phenotype associations were derived from various experimental approaches: RNA interference and genetic allele variations. A gene linked to a particular phenotype is inherently associated with all broader phenotypes that subsume it within the ontology (Supplement, Figs. S1-S2).

### Developmental and Phenotypic spaces

We define a *developmental* space based on the identification of cell types across all quantified cells in the dataset. These types are inferred using gene markers and span different embryo stages. Each gene is thus represented as a “developmental” vector, 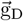, indicating the proportion of cells corresponding to a specific coordinate (cell type × embryonic stage) in which the gene is expressed (Figure 1A; further details can be found in Supplement, Fig. S6). To define the *phenotypic* space we create the gene-phenotype association matrix. Matrix rows indicate the presence or absence of phenotypes associated with the alteration of a specific gene. Given that ontologies often contain redundant structures, we approximate this matrix using NMF (Lee and Seung [1999]). The resulting phenotypic vector for each gene, 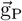, represents the weights of that gene across the dimensions of the feature matrix, W (Figure 1A; further details can be found in Supplement –Figs. S3-S4–, which also discusses alternative definitions of these spaces, Fig. S8).

### Pairwise similarities

We obtain the similarity (*sim*_D_, and *sim*_P_) between pairs of genes as the cosine distance, 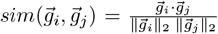. Cosine similarity is used in gene expression analysis when the focus is on expression patterns rather than the magnitude of expression. Two proportional vectors will have a similarity of 1.

### Developmental and phenotypic patterns for D-P, D-p, and d-P gene classes

For developmental patterns, we calculate the fraction of cell types expressing each gene at each embryonic stage. We then compute the average profile for the genes in each class (`D-P`, etc.). For phenotypic patterns, we average the NMF phenotypic profiles of individual genes within each class.

### Associations between specific cell types and NMF phenotypes

We analyze the relationship between the fraction of total genes expressed by each single cell and their relative contribution to a specific NMF phenotype. By the latter we mean the sum of the weights in the NMF matrix *W* corresponding to genes expressed in the individual cell and associated with that specific NMF phenotype. Then, we perform a regression analysis using data from all individual cells, deriving a regression line that captures the overall trend for the given NMF phenotype. The deviations of each cell from this trend are quantified as residuals, calculated as the differences between observed values and those predicted by the regression line. Grouping the cells into subsets belonging to the same cell type, we perform a KS test of the residual distributions of these subsets versus the rest of the cells. Significant (*p*-*value*<0.0001) and positive KS statistics indicate that the specific NMF phenotype is related to that cell type (recall that this statistic is bounded between 0 –perfect agreement between distributions–, and 1 –maximally different). We perform this procedure for all the NMF phenotypes (100 components). For each analyzed NMF phenotype, we included only cell types with more than 50 labeled cells (137 cell types).

### Kolmogorov-Smirnov test

Kolmogorov-Smirnov test compares the sample cumulative distribution *F* (*x*) against the reference cumulative distribution *G*(*x*). The distributions do not need to be parametric. For this we use the `kstest` function of `scipy.stats`, with the parameter ‘greater’ (one sided test). In this case, the null hypothesis is that *F* (*x*) <= *G*(*x*) for all *x*; the alternative is that *F* (*x*) > *G*(*x*) for at least one *x*.

### Gene Ontology enrichment analysis

We downloaded Gene Ontology (GO) annotations from Worm-Base (version WS294). We built three gene–GO term association matrices one for each aspect of the ontology (molecular function, cell component and biological process), by expanding direct annotations to ancestor terms in the GO hierarchy. We calculated the enrichment of terms for different subsets of genes identified along the analysis (Fisher’s exact test; fisher exact function from `scipy.stats`). The test accounts for the total number of genes in both the entire dataset and each subset. We corrected p-values within each GO aspect for multiple testing using False Discovery Rate (FDR).

## Acknowledgements

This work was supported by grants PID2022-140017OB-C22 (MC), PID2019-106116RB-I00 (JFP), and PID2023-151289NB-I00 (JFP) funded by MICIU/AEI/10.13039/501100011033 and by “ERDF/EU” and Grant PRE2021-099926 funded by MICIU/AEI /10.13039/501100011033 and by FSE+ (AL).

## Code availability

Github: https://github.com/ali4lou/D-P-foundations

Zenodo: https://zenodo.org/records/14629057

## References

Pere Alberch. Ontogenesis and Morphological Diversification. American Zoologist, 20(4):653–667, November 1980. doi: 10.1093/icb/20.4.653.

Carmen Bravo González-Blas, Seppe De Winter, Gert Hulselmans, Nikolai Hecker, Irina Matetovici, Valerie Christiaens, Suresh Poovathingal, Jasper Wouters, Sara Aibar, and Stein Aerts. SCENIC+: single-cell multiomic inference of enhancers and gene regulatory networks. Nature Methods, 20(9):1355–1367, September 2023. doi: 10.1038/s41592-023-01938-4.

James A. Briggs, Caleb Weinreb, Daniel E. Wagner, Sean Megason, Leonid Peshkin, Marc W. Kirschner, and Allon M. Klein. The dynamics of gene expression in vertebrate embryogenesis at single-cell resolution. Science, 360(6392):eaar5780, June 2018. doi: 10.1126/science.aar5780.

Diego Calderon, Ronnie Blecher-Gonen, Xingfan Huang, Stefano Secchia, James Kentro, Riza M. Daza, Beth Martin, Alessandro Dulja, Christoph Schaub, Cole Trapnell, Erica Larschan, Kate M. O’Connor-Giles, Eileen E. M. Furlong, and Jay Shendure. The continuum of Drosophila embryonic development at single-cell resolution. Science, 377(6606):eabn5800, August 2022. doi: 10.1126/science.abn5800.

Sean B. Carroll, Jennifer K. Grenier, and Scott D. Weatherbee. From DNA to diversity: molecular genetics and the evolution of animal design. Blackwell publ, Malden (Mass.), 2nd ed edition, 2005.

Silvia Domcke and Jay Shendure. A reference cell tree will serve science better than a reference cell atlas. Cell, 186(6):1103–1114, March 2023. doi: 10.1016/j.cell.2023.02.016.

Gerald M. Edelman and Joseph A. Gally. Degeneracy and complexity in biological systems. Proceedings of the National Academy of Sciences, 98(24):13763–13768, November 2001. doi: 10.1073/pnas.231499798.

Rebecca A. Green, Renat N. Khaliullin, Zhiling Zhao, Stacy D. Ochoa, Jeffrey M. Hendel, Tiffany-Lynn Chow, HongKee Moon, Ronald J. Biggs, Arshad Desai, and Karen Oegema. Automated profiling of gene function during embryonic development. Cell, 187(12):3141–3160.e23, June 2024. doi: 10.1016/j.cell.2024.04.012.

Tamar Hashimshony, Martin Feder, Michal Levin, Brian K. Hall, and Itai Yanai. Spatiotemporal transcriptomics reveals the evolutionary history of the endoderm germ layer. Nature, 519(7542):219–222, March 2015. doi: 10.1038/nature13996.

Chelsea E. Lee, Kaela S. Singleton, Melissa Wallin, and Victor Faundez. Rare Genetic Diseases: Nature’s Experiments on Human Development. iScience, 23(5):101123, May 2020. ISSN 25890042. doi: 10.1016/j.isci.2020.101123.

Daniel D. Lee and H. Sebastian Seung. Learning the parts of objects by non-negative matrix factorization. Nature, 401(6755):788–791, October 1999. doi: 10.1038/44565.

Jun Liu and John Isaac Murray. Mechanisms of lineage specification in Caenorhabditis elegans. GENETICS, 225(4):iyad174, December 2023. doi: 10.1093/genetics/iyad174.

Longqi Liu, Ao Chen, Yuxiang Li, Jan Mulder, Holger Heyn, and Xun Xu. Spatiotemporal omics for biology and medicine. Cell, 187(17):4488–4519, August 2024. doi: 10.1016/j.cell.2024.07.040.

Jonathan S. Packer, Qin Zhu, Chau Huynh, Priya Sivaramakrishnan, Elicia Preston, Hannah Dueck, Derek Stefanik, Kai Tan, Cole Trapnell, Junhyong Kim, Robert H. Waterston, and John I. Murray. A lineage-resolved molecular atlas of C. elegans embryogenesis at single-cell resolution. Science, 365 (6459):eaax1971, September 2019. doi: 10.1126/science.aax1971.

Chengxiang Qiu, Beth K. Martin, Ian C. Welsh, Riza M. Daza, Truc-Mai Le, Xingfan Huang, Eva K. Nichols, Megan L. Taylor, Olivia Fulton, Diana R. O’Day, Anne Roshella Gomes, Saskia Ilcisin, Sanjay Srivatsan, Xinxian Deng, Christine M. Disteche, William Stafford Noble, Nobuhiko Hamazaki, Cecilia B. Moens, David Kimelman, Junyue Cao, Alexander F. Schier, Malte Spielmann, Stephen A. Murray, Cole Trapnell, and Jay Shendure. A single-cell time-lapse of mouse prenatal development from gastrula to birth. Nature, 626(8001):1084–1093, February 2024. doi: 10.1038/s41586-024-07069-w.

Gary Schindelman, Jolene S Fernandes, Carol A Bastiani, Karen Yook, and Paul W Sternberg. Worm Phenotype Ontology: Integrating phenotype data within and beyond the C. elegans community. BMC Bioinformatics, 12(1):32, December 2011. doi: 10.1186/1471-2105-12-32.

Daniel E. Wagner, Caleb Weinreb, Zach M. Collins, James A. Briggs, Sean G. Megason, and Allon M. Klein. Single-cell mapping of gene expression landscapes and lineage in the zebrafish embryo. Science, 360(6392):981–987, June 2018. doi: 10.1126/science.aar4362.

Momoko Watanabe, Jessie E. Buth, Neda Vishlaghi, Luis De La Torre-Ubieta, Jiannis Taxidis, Baljit S. Khakh, Giovanni Coppola, Caroline A. Pearson, Ken Yamauchi, Danyang Gong, Xinghong Dai, Robert Damoiseaux, Roghiyh Aliyari, Simone Liebscher, Katja Schenke-Layland, Christine Caneda, Eric J. Huang, Ye Zhang, Genhong Cheng, Daniel H. Geschwind, Peyman Golshani, Ren Sun, and Bennett G. Novitch. Self-Organized Cerebral Organoids with Human-Specific Features Predict Effective Drugs to Combat Zika Virus Infection. Cell Reports, 21(2):517–532, October 2017. doi: 10.1016/j.celrep.2017.09.047.

Mary Jane West-Eberhard. Developmental plasticity and evolution. Oxford University Press, Oxford New York, 2003.

Long Xiao, Duchangjiang Fan, Huan Qi, Yulin Cong, and Zhuo Du. Defect-buffering cellular plasticity increases robustness of metazoan embryogenesis. Cell Systems, 13(8):615–630.e9, August 2022. doi: 10.1016/j.cels.2022.07.001.

Jianzhi Zhang. Patterns and Evolutionary Consequences of Pleiotropy. Annual Review of Ecology, Evolution, and Systematics, 54(1):1–19, November 2023. doi: 10.1146/annurev-ecolsys-022323-083451.

Lihua Zou, Sira Sriswasdi, Brian Ross, Patrycja V. Missiuro, Jun Liu, and Hui Ge. Systematic Analysis of Pleiotropy in C. elegans Early Embryogenesis. PLoS Computational Biology, 4(2):e1000003, February 2008. doi: 10.1371/journal.pcbi.1000003.

