## Supplement for "Cell atlases and the Developmental Foundations of the Phenotype"

### Extended analysis of phenotypic and developmental spaces

#### Phenotypic space

The Worm Phenotype Ontology (WPO) is an ontology of nematode phenotypes, representing relationships between a controlled vocabulary of phenotypic terms (Schindelman et al. [2011]). Terms are organized as a directed acyclic graph, where a term can be linked to several parental phenotypes. Figure S1 shows the ancestor terms of the phenotype ‘G1 checkpoint variant’.

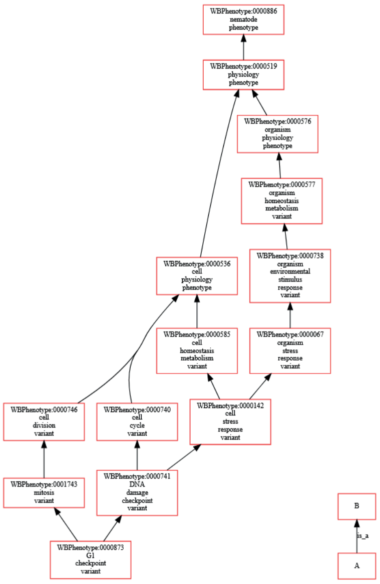


Figure S1: **Representation of the phenotypic space as a directed acyclic graph.** This example illustrates the ancestors of the ‘G1 checkpoint variant’ (a specific term with no children), up to ‘nematode phenotype’ (the root term in the ontology).

##### Gene-phenotype associations

We downloaded gene-phenotype associations from the WormBase (`http://www.wormbase.org/`, version WS290; 10,407 genes linked to phenotypes and 119,901 associations). Most genes have a small number of phenotype annotations (Fig. S2A). All the associations correspond to genetic perturbations: 74% of them correspond to RNA interference (RNAi) studies and the rest (26%) to specific mutations (genetic allele variations) (Fig. S2B).

##### Gene profiles in the phenotypic space

To map genes onto the multidimensional space of phenotypes, we first constructed a matrix of associations between genes and phenotypes, denoted as $V$ (genes $\times$ phenotypes, Fig. S3A). When a gene is annotated with a phenotype in WormBase, we set the corresponding coordinate in $V$ to 1, as well as the coordinates for all its ancestor terms in the ontology.

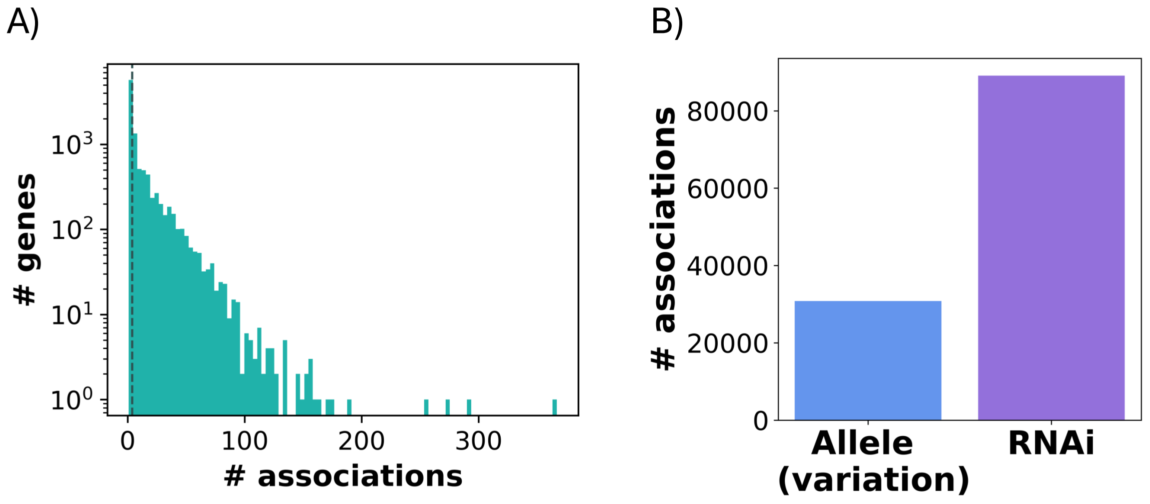


Figure S2: **Gene-phenotype associations.** A) Distribution of the phenotype associations per gene. The dashed line indicates the median. B) Distribution of allele variation and RNA interference (RNAi) perturbations linked to phenotype associations.

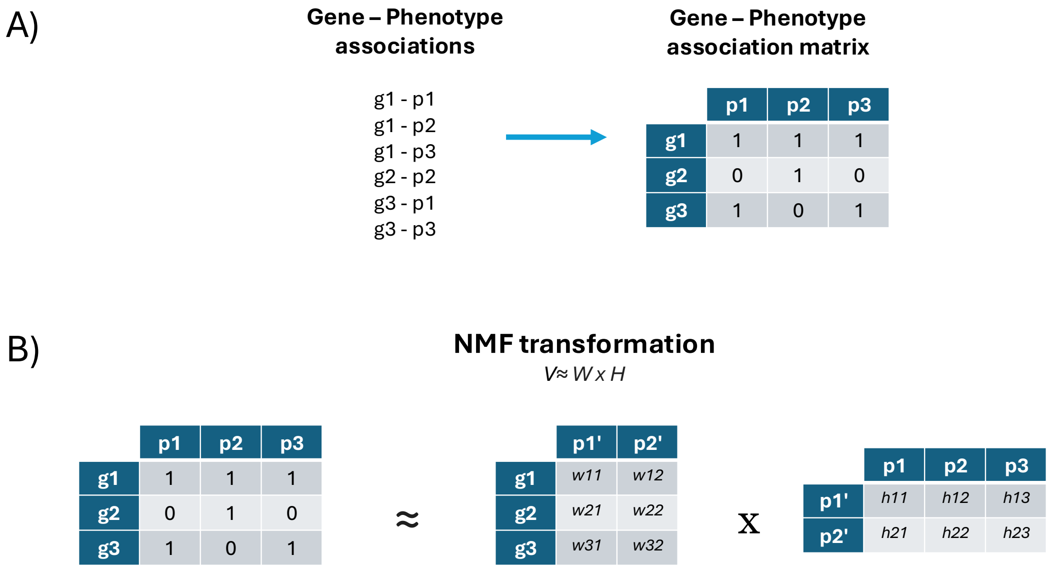


Figure S3: **Gene phenotypic profiles.** A) Using gene-phenotype association data, we construct the gene-phenotype association matrix ($V$), which represents genes within the complete phenotypic space. B) Applying non-negative matrix factorization (NMF) to the gene-phenotype association matrix ($V$) produces two matrices: $W$ and $H$.

##### Non Negative Matrix Factorization (NMF)

To reduce the dimensionality of the phenotypic space, while also reducing the redundancy of the phenotypes, we decomposed the $V$ matrix (genes x phenotypes) by means of non-negative matrix factorization (NMF, Lee and Seung [1999]), where $V \approx W \times H$ (Fig. S3B). $W$ represents genes in the reduced phenotypic space, and H captures the relationships between the new phenotypic components and the original phenotypes.

To choose the number of new phenotypic components in the reduced space, we calculated the mean squared error of the reconstructed $W \times H$ matrix with respect to the initial phenotype matrix $V$ (Fig. S4A; we selected 100 new components based on the curve elbow). Moreover, to check the independence of the new components we computed the correlation between them using the $H$ matrix. The correlation between components is low (Fig. S4B). Figure S4C quantifies their stability.

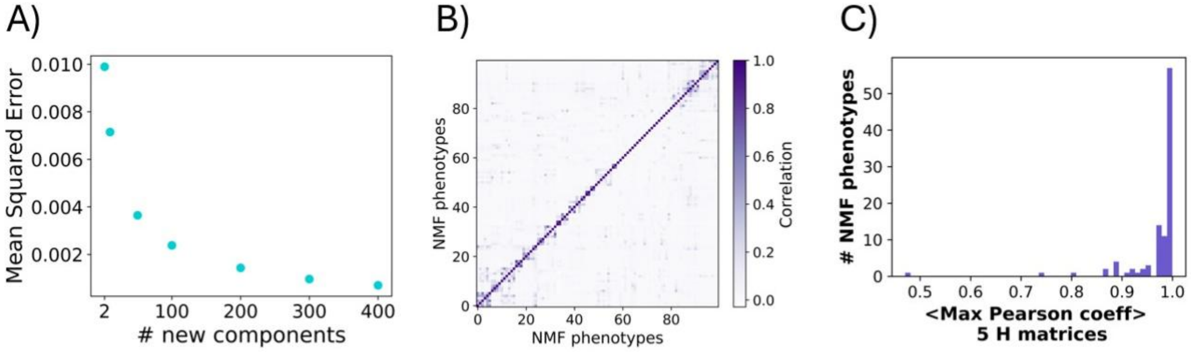


Figure S4: **NMF decomposition.** A) Mean squared error with respect to the number of new phenotypic components. B) Correlation between the new components ($H$ matrix). C) Distribution of NMF component stability. The figure shows the stability of NMF components, measured by comparing the reference $H$ matrix to five new $H$ matrices generated by rerunning the NMF algorithm (NMF involves an optimization, and this process is typically initialized with random values). For each component in the reference $H$, the highest Pearson correlation with components in the new $H$ matrices was identified and averaged across the five runs. The resulting stability values are shown as a distribution.

#### Developmental space

We built the development space from the single cell RNA sequencing data from (Packer et al. [2019]). Specifically, we downloaded the unique molecular identifier (UMI) count matrix, where rows represent individual cells and columns correspond to genes (89,701 cells and 20,222 genes). Figure S5A shows the distribution of the total UMI counts per cell. The maximum UMIs in a cell is 126184, and the minimum is 484. The median = 1,557 UMIs. Figure S5B shows the distribution of expressed genes per cell. The maximum number of cells expressing a gene is 82,912, and the minimum is 0. The median is 543 cells. To filter out empty droplets we discarded cells with <1000 UMIs. We also filtered out genes that were expressed in <3 cells. The shape of the filtered matrix is: 69,612 cells × 18,234 genes.

Each cell has an associated estimated embryo time. Most cells were annotated with a cell lineage, a cell type and/or a cell subtype by the authors (Fig. S5C). Cells with only lineage labels correspond to earlier embryo times (Fig. S5D). We relabeled each individual cell with our own 'cell type' label (229 distinct labels). To construct those labels we took all the possible combinations between the Packer *et al.* cell types and cell subtypes, including 'nan' labels for those cells that do not contain a cell type or cell subtype.

We included in our analysis only those cells with identified cell type or subtype information, resulting in a total of 49,049 cells. While the complete lineage of *C. elegans* is well characterized due to its relatively small and invariant number of cells, this level of detail is impractical to achieve in organisms with significantly larger and more variable cell populations. In such cases, the use of cell type or sub-type labeling becomes essential for categorizing cellular identities and functions. This labeling not only facilitates comparative analyses within a given organism but also provides a framework for extrapolating findings across species, bridging the gap between well-characterized model organisms and more complex systems.

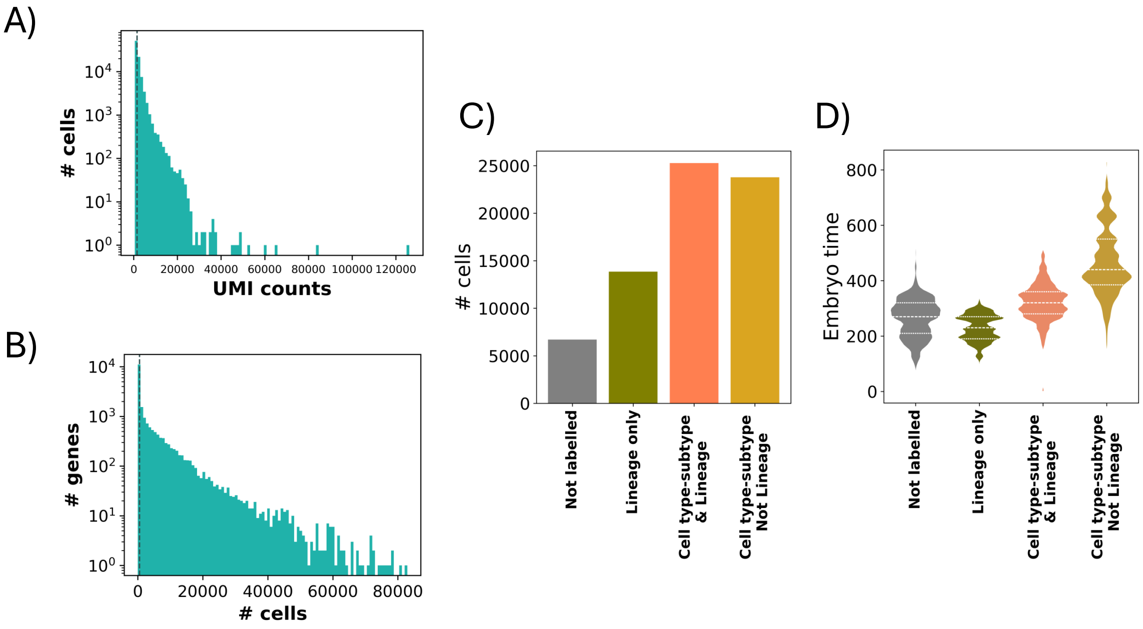


Figure S5: **UMI count matrix statistics.** A) Distribution of total UMIs per individual cell. B) Distribution of total cells expressing a specific gene. In A) and B) dashed lines represent median values. C) Number of cells with different types of Packer *et al.* labels. D) Distribution of embryo times associated with those labels.

##### Definition of the developmental space

We first define a developmental space with two dimensions: embryo time and cell type. Each coordinate of this developmental space will be associated with a specific embryo time ($t_i$), with $t_i$ up to 830 min after first cleavage, and one of 229 possible cell types. The dimension of this space is 136 time points $\times$ 229 cell types. Figure S6A shows the number of cells in each developmental coordinate. Figures S6BC show the number of cells in each embryo time and the number of cells in each cell type, respectively.

##### Gene expression profiles in the developmental space

To represent the expression profile of a gene, we reshaped this developmental space (2D) in a 1-D vector. Each developmental coordinate represents the fraction of cells with time $i$ and cell type $j$, where the gene is expressed.

#### Pairwise similarities

We analyzed the shared genes between the developmental and phenotypic spaces ($n = 8{,}233$ genes). Each gene is represented with two vectors (or profiles), one corresponding the developmental space ($\vec{g}_D$) and

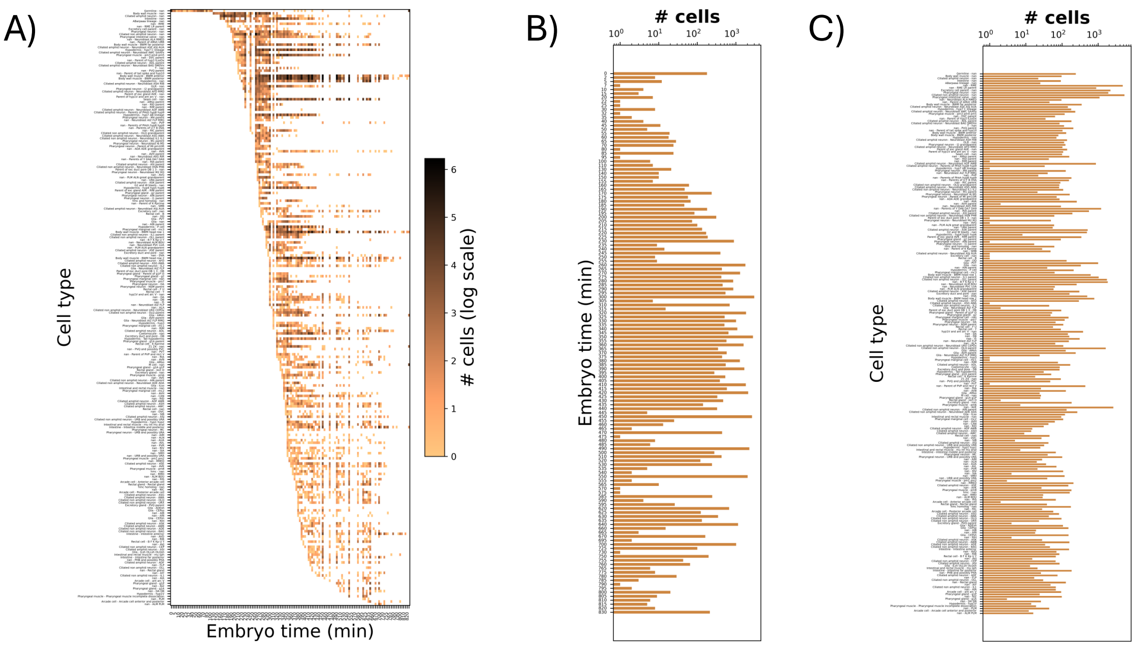


Figure S6: **Developmental space.** A) Number of individual cells sampled at each developmental coordinate (embryo time × cell type). B) Number of cells sampled at each embryo time. C) Number of cells sampled for each cell type.

the other with the phenotypic space ($\vec{g}_{\mathrm{P}}$) (see also main text). We computed all the pairwise similarities between genes in the two spaces using the cosine similarity measure obtaining $sim_{\mathrm{D}}$ and $sim_{\mathrm{P}}$ (Methods, main text).

We then calculated the median similarity of each gene with all the rest of genes in both spaces $\langle sim_{\mathrm{D}} \rangle$ and $\langle sim_{\mathrm{P}} \rangle$ (Fig. S7; Table S2). In Figure S7B we find three repeated similarities: 0.12 (338 genes), 0.14 (287 genes) and 0.07 (258 genes) which correspond to three groups of genes with very similar NMF phenotypic profiles. The first group is characterized by the 97th NMF component (lethal, organism development variant). The second one, by component 6 (embryonic development variant, embryonic lethal), and the third, by components 13 (dauer metabolism phenotype, dauer lifespan variant) and 59 (organismal phenotype, anatomical phenotype).

The average developmental similarity is associated with the number of non-zero developmental coordinates. High $\langle sim_{\mathrm{D}} \rangle$ corresponds to more ubiquitously expressed genes (Fig. S7C).

#### $\mathcal{D}$–$\mathcal{P}$ rule and deviations using alternative representations

We also examined two binary representations of the developmental and phenotypic spaces as alternative representation. For the development space, we represented whether a gene was expressed in at least one cell at each developmental coordinate time $i$ and cell type $j$. For the phenotypic space, we used the $V$ matrix (see Section 1). We then computed the similarity of all pairs of genes in each binary space using the Jaccard similarity, Eq. (1).

$$sim_J(g_i, g_j) = \frac{g_i \cap g_j}{g_i \cup g_j}. \tag{1}$$

For each gene, we calculated the median similarity with all the rest of genes in both spaces $\langle sim_{\mathrm{D}}^{\mathrm{Jaccard}} \rangle$

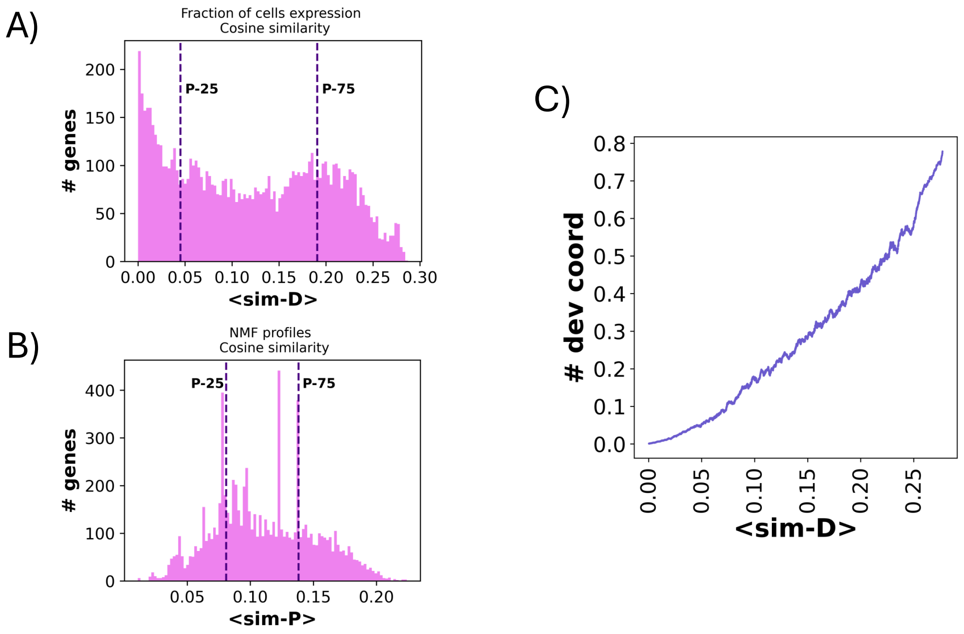


Figure S7: **Median similarities analysis.** A) Distribution of $\langle sim_{\mathrm{D}} \rangle$. B) Distribution of $\langle sim_{\mathrm{P}} \rangle$. C) Number of non-zero developmental coordinates *vs.* $\langle sim_{\mathrm{D}} \rangle$.

and $\langle sim_{\mathrm{P}}^{\mathrm{Jaccard}} \rangle$ (distributions in Figs. S8AB). The $\mathcal{D}$–$\mathcal{P}$ rule holds true, regardless of how developmental and phenotypic expression is quantified. Figures S8CD illustrate the $\mathcal{D}$–$\mathcal{P}$ rule using binary expression profiles from both developmental stages and phenotypic traits.

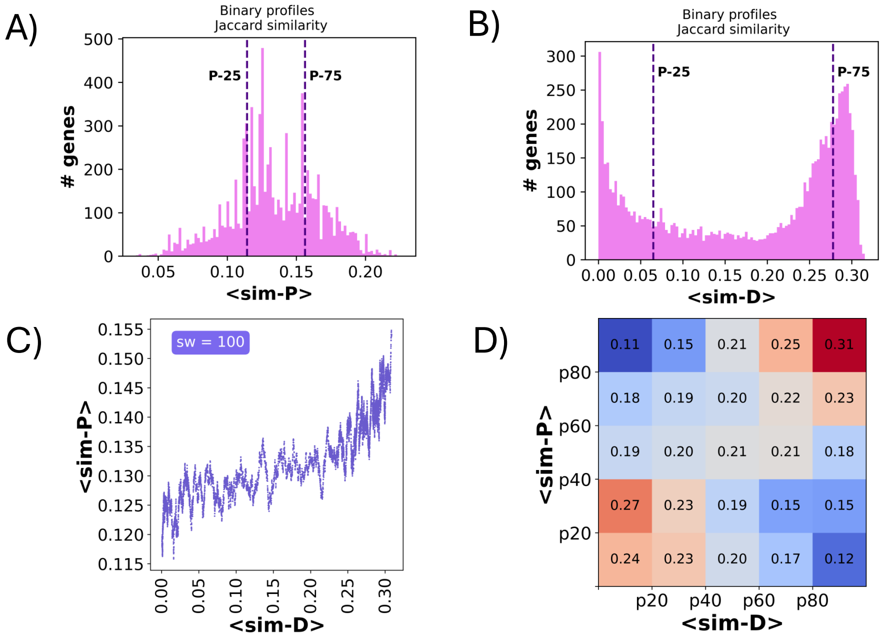


Figure S8: **Binary profiles analysis.** We use Jaccard similarity. A) Distribution of $\langle sim_{\mathrm{D}}^{Jaccard} \rangle$. B) Distribution of $\langle sim_{\mathrm{P}}^{Jaccard} \rangle$. C) $\mathcal{D}$–$\mathcal{P}$ rule confirmation. D) Deviations of the $\mathcal{D}$–$\mathcal{P}$ rule.

#### Deviations of the $\mathcal{D}$–$\mathcal{P}$ rule

We calculated a Loess regression of $\langle sim_{\mathrm{D}} \rangle$ *vs.* $\langle sim_{\mathrm{P}} \rangle$ to study deviations from the $\mathcal{D}$–$\mathcal{P}$ rule (Fig. S9A). We then computed the residuals (distance to the regression fit) for each gene. Based on these residuals, we categorized genes into three groups: `D-P`, `D-p`, and `d-P` genes.

- `D-P` genes (blue, $n$ = 1,322) adhere to the $\mathcal{D}$–$\mathcal{P}$ rule, with $\langle sim_{\mathrm{D}} \rangle$ values higher than the 75th percentile of the $\langle sim_{\mathrm{D}} \rangle$ distribution and residuals that do not deviate by more than one standard deviation ($\sigma$) from the mean of the residual distribution.
- `D-p` genes (purple, $n$ = 168) exhibit low average phenotypic similarity, characterized by residual values below the 10th percentile of the residual distribution and $\langle sim_{\mathrm{D}} \rangle$ values higher than the 75th percentile of its corresponding distribution.
- `d-P` genes (pink, $n$ = 245) show low developmental similarity, characterized by $\langle sim_{\mathrm{D}} \rangle$ values below the 25th percentile of the $\langle sim_{\mathrm{D}} \rangle$ distribution and residuals exceeding the 90th percentile of the residual distribution, indicating high phenotypic similarity.

In Figure S9B, we plot the distributions of the non-zero developmental coordinates: `d-P` genes are expressed in few developmental coordinates (specifically expressed genes), while `D-p` and `D-P` genes are expressed in a high number of developmental coordinates (ubiquitously expressed genes).

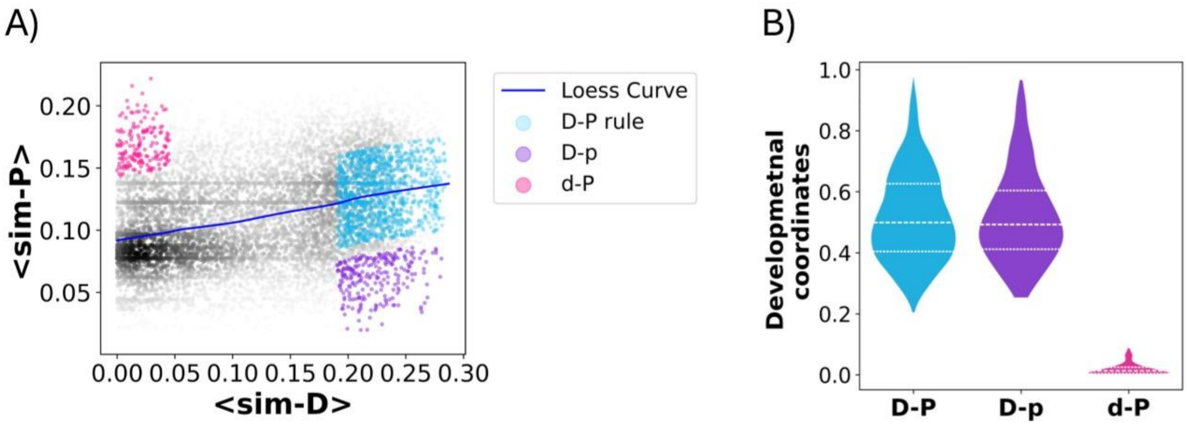


Figure S9: **`D-P` rule deviations.** A) $\langle sim_{\mathrm{D}} \rangle$ *vs.* $\langle sim_{\mathrm{P}} \rangle$ per gene. The blue line is the Loess curve. We define three sets of genes to analyze: `D-P` genes (blue), `D-p` genes (purple) and `d-P` genes (pink). B) Distributions of non-zero developmental coordinates.

We derived the *typical* developmental profile and phenotypic profile for each group of genes (`D-P`, `D-p` and `d-P`) (Fig. 2, main text). For the (typical) developmental profile, we first calculated the fraction of cell types expressing a given gene within each embryo time. We then averaged (mean value) these expression profiles considering the set of genes of a given class (`D-P`, etc.). To obtain the (typical) phenotypic profile, we averaged the NMF phenotypic profiles of the genes within each group.

To illustrate deviations from the `D-P` rule, in Figure S10, we compared the profiles of `D-p` and `d-P` (y-axis) against the `D-P` profile (x-axis). These comparisons are based on the typical profiles shown in

Figure 2, main text. Specifically, each point in the scatter plots represents the values from these typical profiles. The D-P profile is plotted against itself as a reference line, serving as a baseline to identify quantitative differences. Points above or below this reference line indicate deviations: points above the line show where the values of D-p or d-P exceed the corresponding values of the D-P profile, while points below the line highlight where the values fall below the reference. These comparisons are shown in both developmental (Figs. S10AB) and phenotypic (Figs. S10CD) contexts, helping to reveal the extent to which the profiles of D-p and d-P deviate from the expected D-P pattern.

As expected, D-p and D-P genes show comparable expression levels at the same embryo times (Pearson's $\rho = 0.996$, $p$-value $= 3.66 \times 10^{-141}$, Fig. S10A). In contrast, d-P genes are only moderately correlated with D-P genes ($\rho = 0.50$, $p$-value $= 6.81 \times 10^{-10}$, Fig. S10B). This is due to the fact that d-P are expressed in a notably smaller number of developmental coordinates (Figure S9B). Some of them are generally expressed at very low levels, while others are typically lowly expressed but show high or moderate expression in specific coordinates.

For the phenotypic profiles, the correlation between d-P and D-P genes ($\rho = 0.76$, $p$-value $= 5.20 \times 10^{-20}$, Fig. S10C) is high, although lower than the conservation of developmental context (D-P *vs.* D-p). This is explained by the fact that D-P and D-p genes show $\langle sim_{\mathrm{D}} \rangle$ within the same range (Fig. S9A), while $\langle sim_{\mathrm{P}} \rangle$ of d-P genes is, on average, higher than that observed in the D-P genes. The phenotypic correlation between D-p and D-P genes is moderate ($\rho = 0.40$, $p$-value $= 2.69 \times 10^{-5}$).

To better understand the differences between these two groups in the phenotypic space, we selected the phenotypic components with a higher value in the D-p profile than in the D-P one (components #96, #72 and #86; Fig. 2, main text). These components are associated with 'neuron development variant' (comp. #96), 'neuron morphology variant' (comp. #72) and 'cytoplasmic appearance defective early embryo' and 'cell-cell contacts abnormal' (comp. #86). Although D-p genes are expressed broadly during development (Fig. S9B), their impact in the phenotype is highly specific, explaining their lower $\langle sim_{\mathrm{P}} \rangle$ values.

Component #87 (associated with 'germ cell cytoplasmic morphology variant') is characteristic of both D-P and D-p groups but not of the d-P group. This is consistent with the observation that D-P and D-p genes are highly expressed during germline development, whereas d-P genes are not.

To complete the analysis, we performed a phenotype enrichment for each group of genes. The list of enriched phenotypes ($p$-value $< 1 \times 10^{-3}$) is provided in Table S4. Enriched phenotypes in the D-P group and d-P groups include 'lethal', 'organism development variant', 'growth variant' and 'sterile'. The enriched phenotypes in the d-P and D-P classes are predominantly systemic and general, reflecting their organism-wide impact. The majority of enriched phenotypes in the d-P group are also enriched in the D-P group.

The D-p group is enriched in 'body wall muscle myosin organization defective', 'neuron migration variant' and 'neurite development variant' which generally refer to more specific phenotypes (muscle and neuron). Enriched phenotypes explain only a small percentage of genes within this group. No overlap is observed between the most enriched phenotypes ($p$-value $< 1 \times 10^{-4}$) of D-p and D-P.

Finally, Gene Ontology (GO) analysis revealed additional differences between the three groups. (Ta-

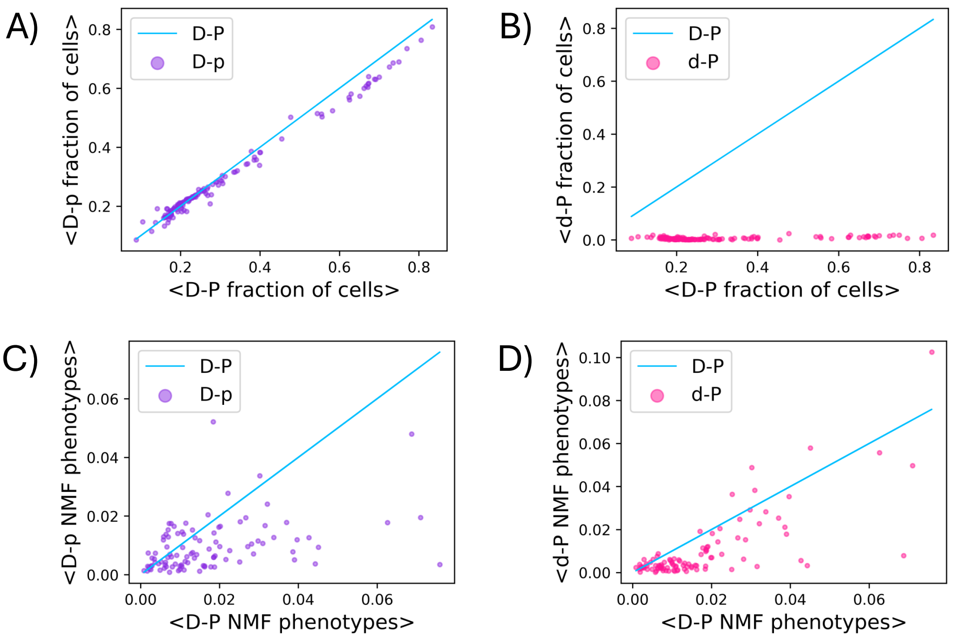


Figure S10: **Comparison of `D-P`, `D-p`, and `d-P` typical profiles in developmental and phenotypic contexts.** In the developmental profile scatter plots (A and B), each point represents the average fraction of cell types expressing genes at each embryonic stage, comparing the subsets `D-p` or `d-P` to `D-P` discussed in Fig. 2 (main text). In the phenotypic context (C and D), each point in the scatter plots corresponds to the average value associated with each NMF phenotype in the profiles of the `D-p` or `d-P` subsets, compared to `D-P`. A) `D-p` *vs.* `D-P` developmental profile values. B) `d-P` *vs.* `D-P` developmental profile values. C) `D-p` *vs.* `D-P` phenotypic profile values. D) `d-P` *vs.* `D-P` phenotypic profile values. The blue line is the representation of the `D-P` profile against itself. Deviations in the average profiles of `D-p` and `d-P` compared to `D-P` are highlighted by points falling above or below this line.

ble S5): **i/**`D-P` genes are enriched in a large number of GO terms, related to housekeeping functions: ‘intracellular organelle’, ‘cytoplasm’ and ‘ribonucleoprotein complex’ (in terms of cellular localization), ‘biosynthetic process’, ‘RNA processing’ and ‘translation’ (in terms of biological processes) and ‘RNA binding’ and ‘structural constituent of ribosome’ (in terms of molecular functions), **ii/** `d-P` genes are enriched in only a small number of GO terms, revealing a general heterogeneity of functions. The enriched terms refer to ‘nucleosome’ and ‘chromatin’, ‘extracellular ligand-gated monoatomic ion channel activity’ and ‘collagen trimer’. Very specific cell functions that are needed for the viability of the organism, and **iii/** `D-p` genes are also enriched in a small number of GO terms, most related to cellular components: ‘cytoplasm’, ‘endomembrane system’, ‘vacuole’ and ‘nucleolus’; and few biological processes ‘rRNA processing’ and ‘protein ufmylation’. Cell components which are present in all the cells but might play specific roles in certain cell types.

### Pleiotropy

The pleiotropic score $\mathbb{P}$ is calculated by adding the scores associated with each gene in the matrix $W$ (obtained by NMF). We defined two classes of genes with extreme $\mathbb{P}$ values based on the pleiotropy distribution (Figure S11): pleiotropic genes ($n$=412, above 95th percentile) and non-pleiotropic genes ($n$=418, below 5th percentile).

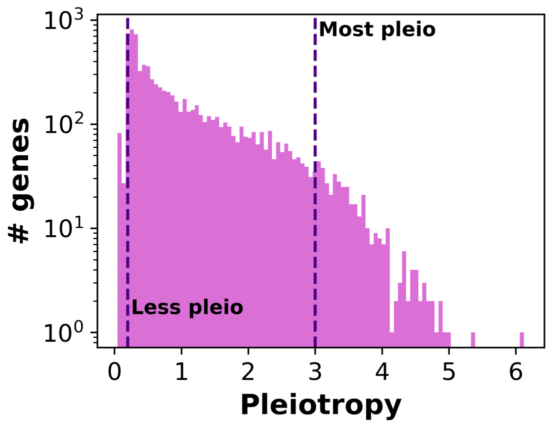


Figure S11: **Pleiotropy distribution.** Dashed lines indicate the 5th and 95th percentiles that define non-pleiotropic and pleiotropic genes, respectively.

#### The biased induced by the hierarchical nature of the ontology

The most specific phenotypes (i.e., those without children) have different depths in the WPO hierarchy (distance from the root term, 'nematode phenotype', Fig. S12A), depending on the branch of the ontology. Thus, genes annotated with deeper terms in the hierarchy will generally be annotated with more phenotypes, leading to an apparent higher pleiotropy value. NMF partially eliminates this bias (Fig. S12B). We compared the pleiotropy ($\mathbb{P}$), based on NMF profiles, with respect to that of the original phenotypic space (defined as the sum of the corresponding row values in the $V$ matrix). Genes associated with specific phenotypes with more than 25 ancestors are represented as orange dots (biased genes). Most of them fall below the linear regression line, indicating that the bias was partially eliminated by NMF.

As NMF is sensitive to initialization, we evaluated the stability of $\mathbb{P}$ across different runs (Fig. S12C). To quantify this, we calculated the coefficient of variation (CV$=\frac{\mu}{\sigma}$), where $\mu$ is the mean and $\sigma$ is the standard deviation of the five $\mathbb{P}$ values for each gene. The CV values remained below 0.3, confirming the consistency in the $\mathbb{P}$ values to initialization variability. Finally, Figure S12D shows the correlation between $\mathbb{P}$ and the number of non-zero components in $W$ (Pearson's $\rho = 0.92$, $p$-value=0.0).

#### Pleiotropic and non-pleiotropic genes

In Figure S13A, we present the distributions of non-zero developmental coordinates where pleiotropic and non-pleiotropic genes are expressed. Most non-pleiotropic genes are expressed in very few developmental coordinates. Figures S13BC represent the fraction of cell types *vs.* embryo times where a gene is expressed. A gene is considered expressed in a cell type or time if at least one corresponding cell shows expression. This can overestimate gene expression, making genes with limited expression in specific developmental coordinates appear broadly expressed across many cell types or stages. Thus, some non-pleiotropic genes are also expressed in a high fraction of cell types and embryo times (Fig. S13C). Most pleiotropic genes are expressed in a high fraction of cell types and embryo times (Fig. S13B).

Next, we compared pleiotropic and non-pleiotropic genes in terms of their enrichment in phenotypes

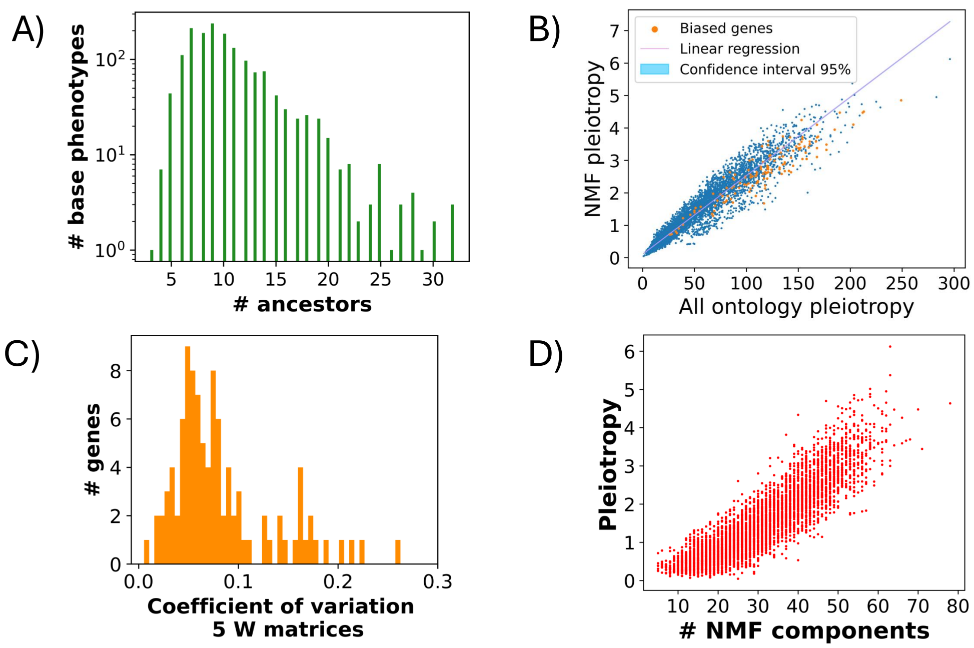


Figure S12: **Redundancy, pleiotropy bias, and pleiotropic score stability.** A) Depth of most specific phenotypes in the WormBase Phenotype Ontology. B) NMF ($W$) pleiotropy vs. $V$ pleiotropy. Orange dots represent those genes associated with at least one of the most specific phenotypes that have more than 25 ancestors in plot A. The major part of those genes fall below the linear regression. C) Coefficient of variation of pleiotropy $\mathbb{P}$ between five NMF runs. D) Pleiotropy $\mathbb{P}$ *vs.* number of non-zero NMF components ($W$ matrix) per gene.

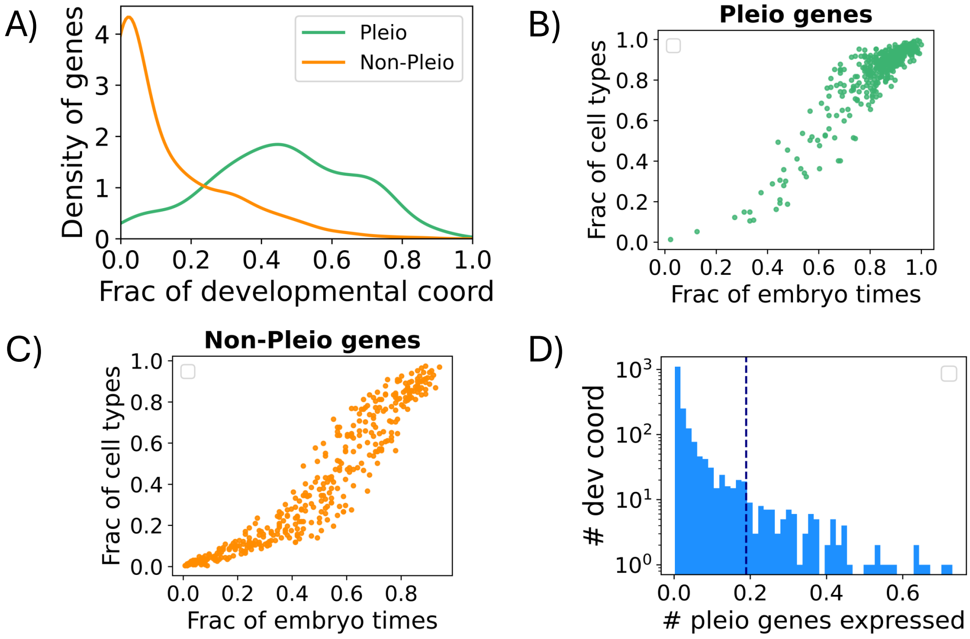


Figure S13: **Pleiotropic and non-pleiotropic genes comparison in development.** A) Distributions of the fraction of non-zero developmental coordinates. B-C) Scatter plot of the fraction of non-zero cell types *vs.* the fraction of non-zero embryo times. B) Non-pleiotropic genes. C)Pleiotropic genes. D) Distribution of the fraction of pleiotropic genes significantly over-expressed in each developmental coordinate. Dashed line marks top 100 values.

(Table S6) and GO terms (Table S7). We find 214 enriched phenotypes for pleiotropic genes. Those with the lowest $p$-value are: 'organ system development variant', 'cell phenotype' and 'organ system morphology variant'. Some more specific enriched phenotypes are: 'cell homeostasis metabolism variant', 'fertility reduced', 'gametogenesis variant', 'gonad morphology variant', 'vulva morphology variant', 'vesicle trafficking variant', 'feeding behavior variant' and 'organism segment morphology variant'. There is just one enriched phenotype for non-pleiotropic genes: 'organism metabolism processing variant'.

With respect to the GO terms, the pleiotropic genes are enriched in a high number of them including: 'intracellular anatomical structure', 'cellular process', 'anatomical structure development', 'cellular component organization or biogenesis' and 'positive regulation of biological process'. Non-pleiotropic genes are enriched in just 6 terms such as 'zinc ion binding' and 'DNA-binding transcription factor activity'. Those terms associate with a small subset of non-pleiotropic genes.

#### Relationship between pleiotropy and development

To examine to what extent pleiotropic genes are particularly expressed at certain developmental coordinates, we first computed the probability of expression of a given gene $g_a$ as the ratio between the total number of cells in which it is expressed, $n_a$, and the total number of sampled cells in the full developmental space, $N$, $p_a = \frac{n_a}{N}$.

Following this, the probability that a gene $g_a$ is expressed in a specific coordinate $(i, j)$ is given by the binomial distribution $P(X = x_{ij}) = \binom{m_{ij}}{x_{ij}} p_a^{x_{ij}} (1 - p_a)^{m_{ij} - x_{ij}}$. This is the probability of having $x_{ij}$ cells expressing the gene in a coordinate $(i, j)$ with a total of $m_{ij}$ sampled cells. Given this distribution, we introduce the $z$-score$(g_a, i, j)$=$\frac{x_{ij} - m_{ij} p_a}{\sqrt{m_{ij} p_a (1-p_a)}}$. We also calculated an over-expression $p$-value by adding the probability of obtaining a number of cells equal or higher than $x_{ij}$ in a coordinate: $p$-value$(g_a, i, j) = P(X \geq x_{ij}) = 1 - P(X < x_{ij}) = 1 - P(X \leq x_{ij} - 1)$. To assess whether a gene is over-expressed at each coordinate, we used the criteria z-score>2 and $p$-value<0.001.

For each coordinate, we computed the fraction of pleiotropic genes over-expressed based on the previous criterion. Figure S13D depicts the distribution of developmental coordinates with a given fraction of over-expressed pleiotropic genes. From this data, we focused on the top 100 coordinates with the highest values (above the dashed line in Figure S13D) for further analysis (Section 3, main text, Fig. 3C). The standout cell types in Fig. 3C of the main text are:

- All the intestine cell types.
- Pharyngeal 1/12: 'Pharyngeal marginal cell - mc1'.
- Body Wall Muscle 4/6: 'Body wall muscle - nan', 'Body wall muscle - BWM anterior', 'Body wall muscle - BWM posterior' and 'Body wall muscle - BWM head row 1'.
- Neuron: (3/140) 'Ciliated amphid neuron - nan', 'Ciliated non amphid neuron - nan', and 'Ciliated amphid neuron - Neuroblast ASE ASJ AUA (precursor)'.
- Excretory cells (1/11): 'Excretory cell - nan'.
- Hypodermis (2/11): 'Hypodermis - hyp7 C lineage' and 'Hypodermis - hyp7 AB lineage'
- Precursor cells: 'Germline - nan', 'Seam cell - nan' and 'M cell - nan'.

Note that 8 of the 20 highlighted cell types could not be annotated with cell subtype and appear with the label 'nan', which could be related to early cell types (partially differentiated, but not fully specific).

Additionally, we identified relationships between specific phenotypes enriched with pleiotropic genes and the highlighted cell types shown in Figure 3C of the main text. For example, 'cell homeostasis metabolism variant' can be associated with Excretory cells (removal of metabolic waste) and Intestine cells (digestion and absorption). Also, 'Hypodermis - hyp7 AB lineage' play an important role in vulva formation ('vulva morphology variant'), as they interact with vulva precursor cells. 'Fertility reduced' can be related to Germline. 'Feeding behavior variant' can be associated with the role of Ciliated Amphid Neurons (detecting food-related chemical cues), Intestinal Cells (Middle, Posterior, Anterior for food processing), Pharyngeal Muscles (pm3, pm4, pm5; crucial for food ingestion) and Body Wall Muscles (BWM Anterior, BWM Posterior; locomotion toward food). 'Organism segment morphology variant' can be related to Hypodermis (hyp7 AB lineage, hyp7 C lineage) and Body Wall Muscles (BWM anterior and posterior) important for segment organization, and Seam Cells, for the maintenance of segment boundaries.

With respect to the Figure 3D, main text, we observed that genes with higher $\langle sim_{\mathrm{P}} \rangle$ tend to have higher pleiotropy values (Fig. S14A). This explains how the `d-P` genes are (in general) more pleiotropic than the `D-p` ones (Fig. S14B). The distributions are significantly different (KS=0.21, p-value=9.36x$10^{-5}$).

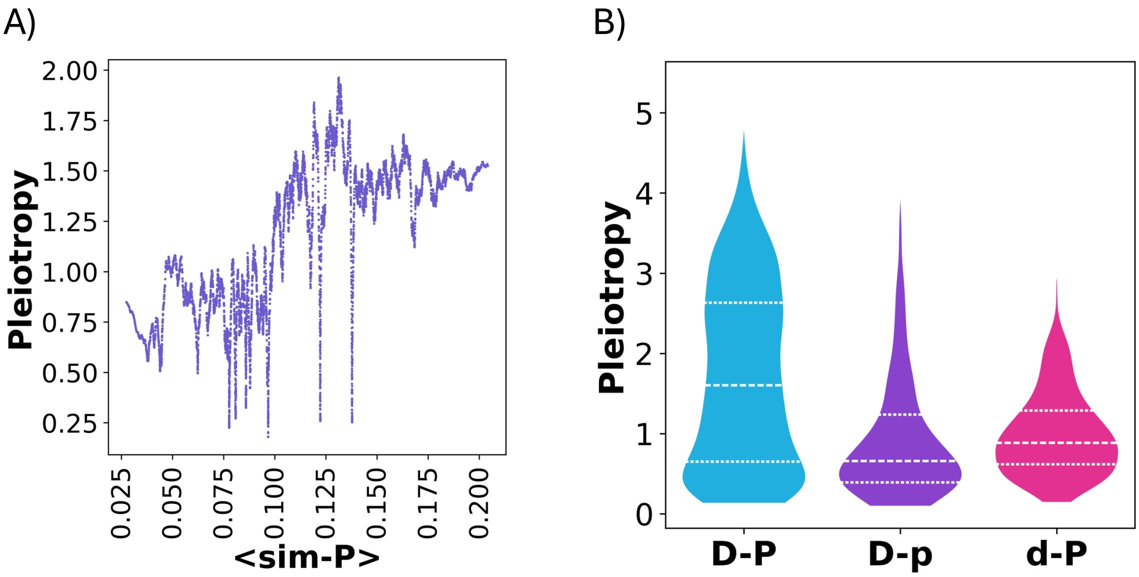


Figure S14: **Relation between pleiotropy and the $\mathcal{D}$–$\mathcal{P}$ rule. A)** Pleiotropy *vs.* $\langle sim_{\mathrm{P}} \rangle$. Sliding window = 100. **B)** Pleiotropy distributions of `D-P`, `D-p` and `d-P` genes.

#### Alternative measures of pleiotropy

We compared our measure of pleiotropy with some earlier work. Zou et al. [2008] provided a list of 19 highly pleiotropic genes. We just found 6 genes from that list in our set of ≈8,000 analyzed genes. 4 of those genes were within *our* pleiotropic genes. Xiao et al. [2022] analyzed a set of 752 gene knockouts but were able to annotate cellular phenotypes for only 331 genes, as detailed in their dataset. Of these 331 genes, 327 overlap with our dataset. We compared $\mathbb{P}$ for these 327 overlapping genes with the number of phenotypes reported in their study for each gene to observe a significant correlation (Spearman's $\rho$=0.57, $p$-value=$3.36\times10^{-25}$; Pearson's $r$=0.53, $p$-value=$3.70\times10^{-22}$).

Finally, Green et al. [2024] analyzed 503 gene knockouts and searched for cellular phenotypes related to the germ layer or morphogenesis. They provided a dataset of associations between those 503 genes and phenotypes IDs from the WormBase (15 IDs related to germ layer and 15 to morphogenesis). The association score was the number of embryos that had that gene knockout and showed the phenotype. We find 475 genes in common with our dataset. We defined *their* pleiotropy value for each gene as the sum of the fractions of embryos in which the knockout of that gene was associated with specific phenotypes. A gene was classified as pleiotropic if its knockout significantly impacted a large proportion of embryos. However, we observed no correlation between that measure of pleiotropy and our $\mathbb{P}$ value.

#### Coarse interpretation of the $\mathcal{D}$–$\mathcal{P}$ rule

The analysis of significant KS statistics allowed us to establish connections between phenotypes and cell types (Methods, main text). For instance, Figure 4B in the main text connects NMF phenotype component #49 with ciliated amphid neurons. Similarly, Figure S15A associates phenotype component #82 with body wall muscle, while Figure S15B links NMF phenotype component #5 with excretory cells.

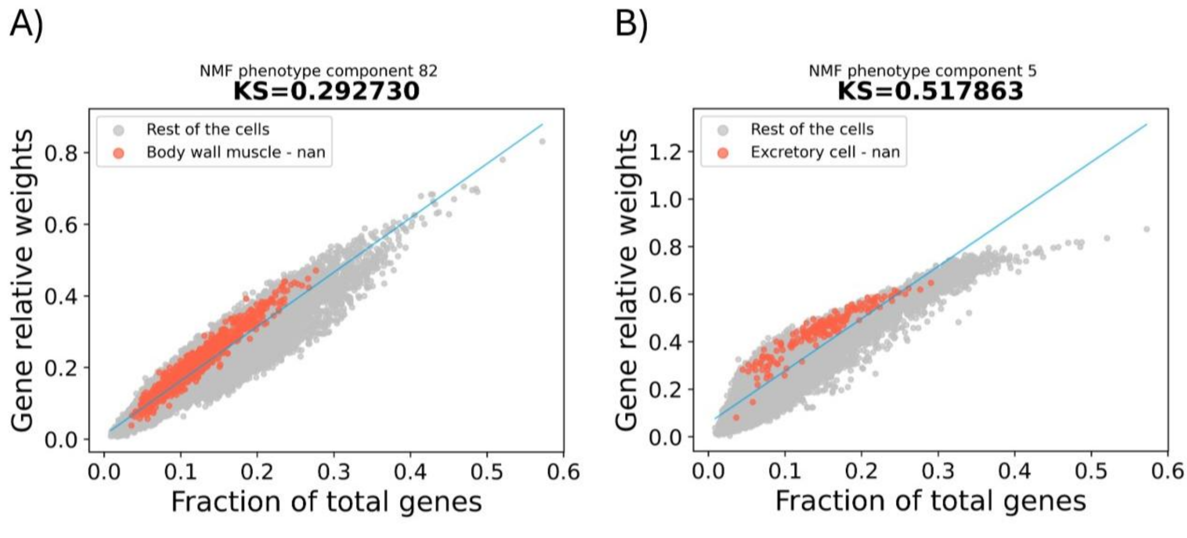


Figure S15: **Associations between a NMF phenotype and a cell type.** Other examples of identifying mediator cells for NMF-derived phenotypes; A) phenotype #82 (body wall muscle cells, KS $\approx$ 0.3) and B) phenotype #5 (excretory cells, KS $\approx$ 0.5).

Fig.S16A illustrates the distribution of the number of cell types mediating phenotypes, ranging from approximately 10 to 70 out of a total of 137 cell types. Fig.S16B presents the inverse perspective, showing how many phenotypes are mediated by each cell type (total of 100 phenotypes). Additionally, note that the higher the KS statistic (ranging from 0 to 1), the stronger the association between a cell type and an NMF-derived phenotype. The distribution of this statistic for all significant cases ($p$-value>0.0001, Fig4C of the main text) is displayed in FigureS16C.

To identify cell types with a high impact on a larger number of phenotypes (and vice versa), we selected associations where the KS statistic exceeded the 75th percentile and counted these for each cell type and NMF phenotype (Table S9). The cell types mediating the greatest number of phenotypes are primarily found in clusters 1 and 5 (Fig. S17A, orange and brown clusters, respectively; this corresponds to the top-down orientation Fig. 4C, main text; see also next section). Within cluster 1, the top cell types are precursor cells, while cluster 5 includes intestinal cells, M cells, glia, and pharyngeal muscle.

Moreover, NMF phenotypes strongly associated with many cell types include those related to chemosensory behavior, organ morphology, cytoskeleton organization, and drug resistance. These phenotypes belong to the red, orange, and purple clusters in the top dendrogram (Fig. 4C, main text). The most relevant cell types are neurons (red and orange clusters) and neuronal precursors (purple cluster), with the abundance of distinct neuron annotations driving more significant nervous system-related associations with high KS values.

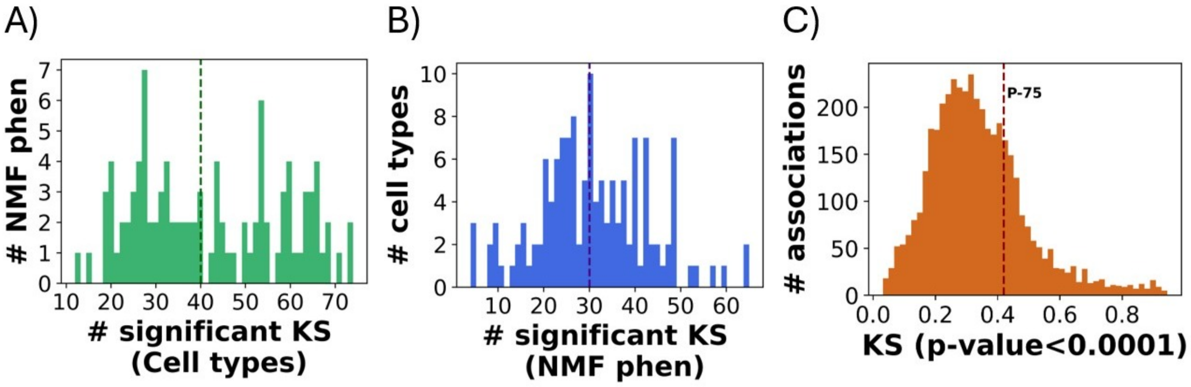


Figure S16: **KS value distributions.** A) Number of significant associated cell types per NMF component. B) Number of significant associated NMF components per cell types. The dashed line in plots A and B represents the median. C) Distribution of all the significant KS values ($p$-value $<$ 0.0001). The dashed line indicates the percentile 75th.

#### Association between lineages and cell types belonging to each cluster

As discused before, the KS association between cell types and NMF components and the posterior clustering allows the cells to be grouped together in more generalized groups that identify the same kind of cell types: such as precursor and parent cell types, cell types related to nervous system and cell types related to different anatomical parts (pharynx, hypodermis, intestine, body wall). These clusters are shown in the dendrogram of Figure 4B of the main text and in Figure S17A.

We also computed the percentage of cells in each cluster associated with each *C. elegans* lineage. By identifying the proportion of each cell type labeled with a lineage and averaging these proportions within clusters, we determined the overall percentage of cells per cluster linked to each lineage. The results are shown in Figure S17B.

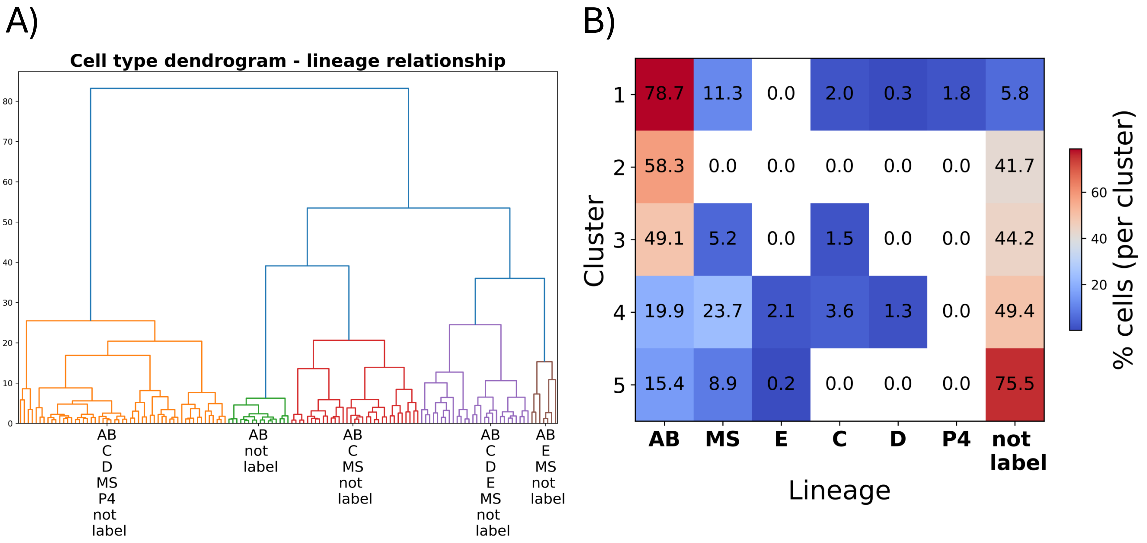


Figure S17: **Association between lineages and cell types' clusters.** A) Dendrogram of the cell types. This dendrogram is obtained from the KS association between NMF components and cell types. It is also shown in Figure 4B of the main text. We find 5 labeled clusters. B) Percentage of cells of each cluster associated with each lineage.

### References

Rebecca A. Green, Renat N. Khaliullin, Zhiling Zhao, Stacy D. Ochoa, Jeffrey M. Hendel, Tiffany-Lynn Chow, HongKee Moon, Ronald J. Biggs, Arshad Desai, and Karen Oegema. Automated profiling of gene function during embryonic development. *Cell*, 187(12):3141–3160.e23, June 2024. doi: 10.1016/j.cell.2024.04.012.

Daniel D. Lee and H. Sebastian Seung. Learning the parts of objects by non-negative matrix factorization. *Nature*, 401(6755):788–791, October 1999. doi: 10.1038/44565.

Jonathan S. Packer, Qin Zhu, Chau Huynh, Priya Sivaramakrishnan, Elicia Preston, Hannah Dueck, Derek Stefanik, Kai Tan, Cole Trapnell, Junhyong Kim, Robert H. Waterston, and John I. Murray. A lineage-resolved molecular atlas of *C. elegans* embryogenesis at single-cell resolution. *Science*, 365 (6459):eaax1971, September 2019. doi: 10.1126/science.aax1971.

Gary Schindelman, Jolene S Fernandes, Carol A Bastiani, Karen Yook, and Paul W Sternberg. Worm Phenotype Ontology: Integrating phenotype data within and beyond the C. elegans community. *BMC Bioinformatics*, 12(1):32, December 2011. doi: 10.1186/1471-2105-12-32.

Long Xiao, Duchangjiang Fan, Huan Qi, Yulin Cong, and Zhuo Du. Defect-buffering cellular plasticity increases robustness of metazoan embryogenesis. *Cell Systems*, 13(8):615–630.e9, August 2022. doi: 10.1016/j.cels.2022.07.001.

Lihua Zou, Sira Sriswasdi, Brian Ross, Patrycja V. Missiuro, Jun Liu, and Hui Ge. Systematic Analysis of Pleiotropy in C. elegans Early Embryogenesis. *PLoS Computational Biology*, 4(2):e1000003, February 2008. doi: 10.1371/journal.pcbi.1000003.
